# Kinference: Pairwise kinship detection for Close-Kin Mark-Recapture

**DOI:** 10.64898/2026.05.18.725841

**Authors:** Mark V. Bravington, Shane M. Baylis, J. Paige Eveson, Pierre Feutry

**Affiliations:** Estimark Research, Hobart, Australia; CSIRO Environment, Hobart, Australia

## Abstract

Close-Kin Mark-Recapture (CKMR) is a statistical framework for estimating demographic parameters of wild populations. Instead of recapturing individuals, it relies on the identification of closely-related pairs such as parents and offspring, or siblings. By measuring how often such close-kin are “recaptured” among sampled animals (whether alive or dead), scientists can estimate demographic parameters such as census size, mortality rates, and connectivity. CKMR is starting to change fisheries and wildlife management by giving more reliable demographic information, even for many species that resist conventional approaches. Here we introduce the `kinference` R package, which provides a set of tools for finding close-kin pairs among thousands of samples each genotyped at thousands of SNPs, and for associated quality control. The CKMR context implies different requirements and assumptions to many other kinship programs. In particular, `kinference` accounts empirically for linkage without requiring a genome assembly, is able to estimate and control false-negative and false-positive probabilities, and can cope with null alleles. The package has been developed and used in numerous CKMR projects since 2017. This paper documents the assumptions, statistical algorithms, and intended workflow for `kinference`.

## 1. Introduction

The software `kinference` is an R package (R Core Team, 2026) that finds pairs of closely-related animals (up to second-order kinship) from genotypes with several thousand SNPs per sample, among datasets comprising hundreds to hundreds-of-thousands of tissue samples. Kin-finding is the core data preparation step for Close-Kin Mark-Recapture (CKMR), described below, and the package is designed specifically with CKMR in mind; thus, it has several features not found in the many other programs that can identify kin. Its origins date back to about 2017, when CSIRO Australia was starting development of large-scale SNP-based CKMR. Since then, various versions of `kinference` have been used in at least ten CKMR projects globally, across fisheries, renewable resource management, and marine and terrestrial conservation. Despite extensive use of `kinference`, until now there has never been a full description of the assumptions and algorithms behind it, covering not just kin-finding *per se* but also some essential Quality Control (QC) steps for loci and samples that should happen after genotypes are called but before kin-finding. As well as technical descriptions, this paper includes insights based on our practical experience with `kinference`. The main notation is shown in Table 1.

**Table 1.**
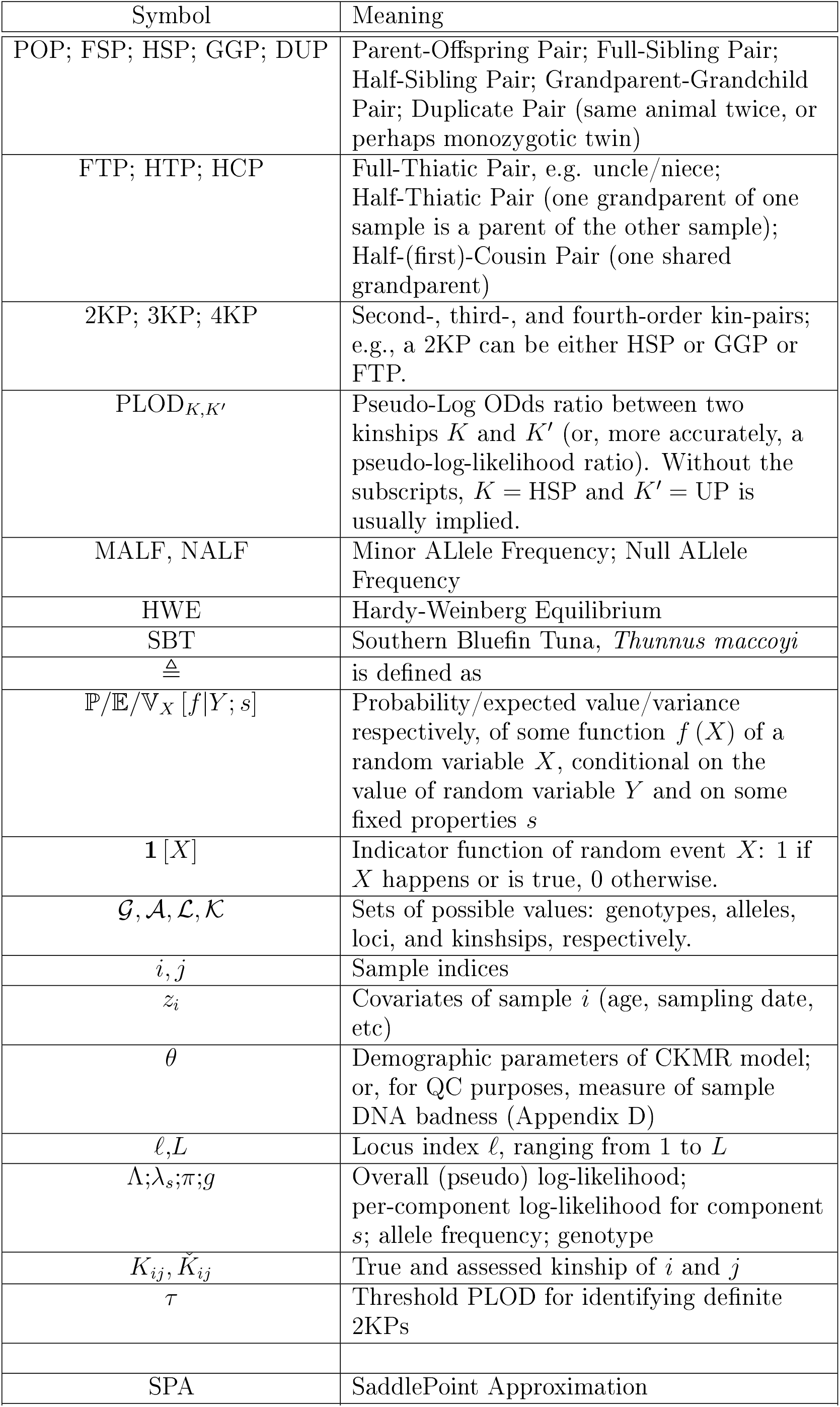
Notation. Only the less-standard usages are shown, and symbols de ned only locally are omitted.

The algorithms in `kinference` are based on well-known principles, albeit with some novel statistical details, and of course kinship and pedigree are long-standing topics of interest in genetics addressed by numerous other programs. Thompson (2000) is a good general reference for the underlying theory. The features that set `kinference` apart are driven by the specific purpose to which it is tailored: CKMR. We brie y review CKMR in section 2, including a description of how kinship uncertainty can be handled efficiently. In section 3, we then summarize the data requirements, assumptions, and outputs of `kinference`, as well as the issue of null alleles in SNPs, which can sometimes be highly important for kin-finding. Sections 4 and 5 describe the kinship computations and the QC statistics respectively (although in practice the QC should be done first). Finally, the discussion in section 6 addresses the statistical desiderata of kin-finding for CKMR versus other purposes, compares `kinference` to other relatedness software, and comments on possible future directions. The Appendices provide additional algebraic details and numerical results. Detailed documentation (e.g. function arguments, le formats, etc.) is mostly left to the `kinference` package (https://closekin.r-universe.dev/kinference), which includes an extensive vignette.

## 2. Close-Kin Mark-Recapture

### 2.1. Close-Kin Mark-Recapture

CKMR, as described in Bravington et al. (2016b), is a framework for estimating population-dynamic parameters of wild animal populations, including adult absolute abundance ( “census population size”) and mortality rates. It relies on pairwise genetic comparisons among large collections of tissue samples. The core idea, which dates back to Skaug (2001), is that every sampled animal had precisely one mother and one father at birth; those two parents are “marked” by the sample’s genotype, and can be “recaptured” either directly (when one sample turns out to be a parent of another) or indirectly (when two samples turn out to share a recent common ancestor, almost always a parent). Intuitively, bigger populations give o spring more freedom to choose their parents, so kin-pairs are rarer. This can be formalized into a statistical framework that embodies situation-specific time-varying population dynamics, individual characteristics such as size and date of capture, and sampling mechanisms, to allow estimation of parameters. A key advantage over conventional mark-recapture is that there is no need for live release: “marking” happened at birth, well before sampling, so that CKMR can be applied to datasets collected lethally, e.g. from hunting as in Taras et al. (2024) (though CKMR can also work with non-lethal biopsies or with a mixture, as in Lloyd-Jones et al., 2023 and Hillary et al., 2018a). CKMR is particularly attractive for fisheries, especially high-value high-problem species like many tunas, because of the direct and low-assumption route to absolute abundance, the comparative simplicity of sampling from catches, and the scalability: sample size requirements are proportional to the square root of abundance (Bravington et al., 2016b, section 4.3), so in general a smaller proportion of catch is needed for larger populations.

The first practical application, which began in 2006 at CSIRO Australia, was to Southern Blue n Tuna (SBT; Bravington et al., 2014). It used 25 microsatellites the only proven option for genetic markers at the time to find Parent-O spring Pairs (POPs) only. From the mid-2010s, with sequencing becoming increasingly reliable and affordable, CKMR projects have generally used SNPs as the markers instead. The ability to genotype several thousand SNPs at comparable cost to (and with much less work than) dozens of microsatellites gave enough power to detect not just POPs but also Half-Sibling Pairs (HSPs), which yield extra demographic information, in particular on adult mortality rate. The largest and most mature CKMR application to date is the SNP version of SBT (Davies et al., 2020b), where by 2025 over 30,000 samples had been genotyped and over 300 POPs and HSPs found, with about 3,000 samples being added per year. The CKMR analysis is a core part of ongoing sampling, stock assessment, and management procedure calculations used in setting triennial catch limits for this near-billion-dollar fishery (Hillary et al., 2018c; Hillary et al., 2018b). SNP-based CKMR has also been used on endangered and/or heavily-depleted species that have resisted assessment with more conventional methods, such as white sharks (Hillary et al., 2018a) and school sharks (Thomson et al., 2020). Most applications have been to sharks and marine scalefish, but there have been recent terrestrial and mammalian applications too (Taras et al., 2024; Lloyd-Jones et al., 2023).

The main kinships of interest are POPs and HSPs, since they convey the most information about recent demographics, and they can be found reliably and affordably (see later). Sometimes it is necessary, and occasionally even useful, to also consider Full-Sibling Pairs (FSPs), Grandparent-Grandchild Pairs (GGPs), and Full-Thiatic Pairs (FTPs, “thiatic” being a genderless term for aunt/nephew-type kinship). Thus, all first-order and second-order kin-pairs (1KPs and 2KPs) are of interest. Third-order pairs (3KPs) and weaker kin have not been used to date, but constitute a significant nuisance in that they cannot completely be distinguished from 2KPs.

There is no “CKMR program” that will set up and fit models automatically; rather, CKMR is a statistical framework for users to write their own models. The basis of any CKMR model is case-specific formulae for the demographic probability of *true* kinship, regardless of genetics. That is, the analyst needs to de ne probabilities of the form ℙ [*K*_*ij*_ = *k*|*z*_*i*_, *z*_*j*_, *θ*], where: *K*_*ij*_ is the true kinship of samples *i* and *j, k* is a kinship of interest, say HSP; *z*_*i*_ and *z*_*j*_ are covariates of *i* and *j*, which vary across applications but usually include sampling date as well as age or some proxy; and *θ* comprises the case-specific demographic parameters that will describe abundance and other phenomena. Not all pairs need be tested for all kinships in the model; depending on covariates and demography, some comparisons are not worth the modeling effort. However, each pair tested for a given kinship is assumed to yield a definitive yes-no outcome, based on genetics and sometimes on covariates, so that *θ* can be estimated by maximizing the sum of pairwise log-likelihoods, or by a Bayesian extension. Sometimes it may also be necessary to allow for uncertainty in the *observable* kinship of a pair, via consideration of false-positive and false-negative rates.

The information content in CKMR comes from the proportion of kin-pairs found per comparison, stratified by target kinship *k* and sample covariates {*z*_*i*_, *z*_*j*_}. Reliable parameter estimates require the number of kin-pairs present to be accurately assessed. The kin-finding task is not just to find “some” kin-pairs of known degree, but to either find all of them or at least to be able to estimate how many have been overlooked: the false-negative rate. In CSIRO’s CKMR projects, depending on the demographic breakdown of sampling and to a lesser extent on the number of loci, the estimated false-negative rate for HSPs using all samples has ranged from practically none to about 40%. Values as high as that are intolerable, but can often be reduced substantially by restricting comparisons to certain covariate-combinations where 3KPs are less likely; false-negative rates within those comparisons actually included in CKMR models have typically been below 20%.

For parameter estimates to be precise (say, no more than a 10% Coefficient of Variation, or CV), the number of POPs and HSPs detected needs to be order-of-magnitude 100 (Bravington et al., 2016b, section 4). In our experience of ecology, it is rare for an abundance estimates of a large population to have a realistic CV below 10%, whether from CKMR or any other method; for better or worse, a 20% CV is often regarded as fairly good. In that context, it is useful to bear in mind that a bias of a couple of percent would not be very important. Models should be “good enough” approximations, but do not need to be perfect.

The sample size required to find enough kin-pairs scales with the square-root of adult abundance (amongst other factors), and in CSIRO projects sample sizes have ranged from roughly a thousand (e.g. Hillary et al., 2018a) to many tens of thousands (e.g. Davies et al., 2020a). The number of pairwise comparisons can therefore be very large (∼ 10^8^ for SBT; ∼ 10^10^ in some proposed projects). Tight control of accuracy when finding kin-pairs is obviously essential when looking for an unknown but small number of “needles” (true kin-pairs) in a very large “haystack” of pairwise comparisons. For large populations, CKMR requires only a small proportion to be sampled, with the consequence that only a small proportion of samples are involved in close-kin pairs, and the number of sampled close-kin triads, etc., is negligible; this is the “sparse sampling” situation described in section 4.3 of Bravington et al. (2016b).

Although this framework for CKMR covers a broad range of population dynamics and sampling scenarios, it does have limits, and they are relevant to the scope of `kinference`. In particular, it would normally be necessary to assume that samples come from a genetically panmictic population (or if not, that probabilistic assignments can be made *a priori* to known underlying populations) and that inbreeding is negligible (and thus that adult census size is not too small). Some further conditions are discussed at the end of Bravington et al. (2016b).

### 2.2. Kinship uncertainty in CKMR

The CKMR framework requires that kin-finding has a definite binary outcome: for example, as far as the model is concerned, two samples are either a POP or they are not. If the genotyping data meets the standards expected by `kinference`, as described shortly, then there should be little or no ambiguity in identifying POPs and FSPs genetically, including distinguishing them from 2KPs and weaker kin. Thus, there is little need to worry about uncertainty for first-order kinships. However, in the absence of genomic information (see Discussion), 2KPs cannot be completely distinguished from 3KPs by genetics, so kinship uncertainty needs somehow to be allowed for. Applications can deal with this by modelling an observable “kinship indicator” 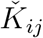 rather than true kinship *K*_*ij*_, where (for example) 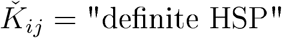 if-and-only-if one or more classification statistics falls into some target range(s). The target range should be chosen by the analyst so that the number of non-HSPs likely to generate a within-range statistic is negligible (see below for comments on HSPs versus other 2KPs). Working with 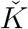 rather than *K* incurs some false-negative probability 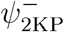 that a true HSP will generate a statistic outside the range. The kinship probability formulae can be adapted as follows:

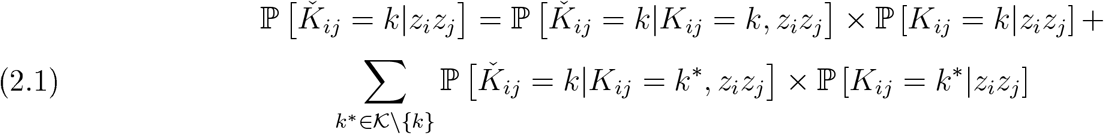

This equation is more subtle than it might appear. The first term in each product, 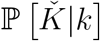, is a purely genetic quantity which depends only on the distribution of the classification statistic amongst pairs with some true particular kinship, and not on demography or the covariates *z*_*i*_ or *z*_*j*_. The second terms, of the form ℙ [*K*|*z*], have nothing to do with genetics but can vary strongly with *z*. Writing 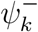 as the false-negative probability and 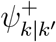 as the false positive probability when the true kinship is *k*′, we have

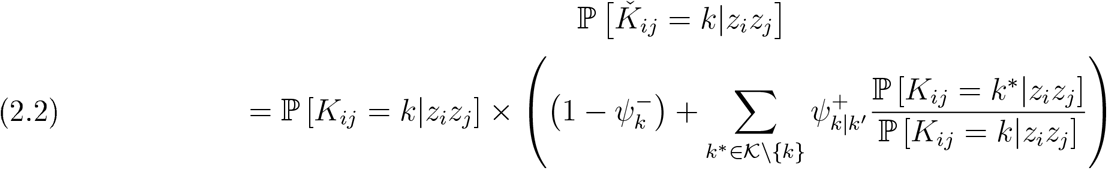

As section 4.2.5 explains, the quantity 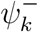 can be estimated empirically given enough true target kin-pairs. The key practical step is to ensure that the right-hand sum, which involves the 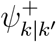 factors, is small relative to 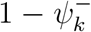 for all useful covariate-strata {*z*_*i*_, *z*_*j*_}: i.e., strata that are demographically informative and might result in an appreciable number of target kin. The issue is not so much the *total* number of false-positives, which might be estimated empirically; but rather that their composition in terms of *z*_*i*_ and *z*_*j*_ could be very different from that of the target kin. For example, if the target kinship is HSP, which will mostly be born in the same generation, then the main risk of confusion is likely to come from HTPs (a type of 3KP), which are typically born one generation apart. Estimates of adult mortality rate are determined basically by the average birth-gap separation of HSPs, so even a small number of contaminating HTPs with much longer birth-gaps could introduce appreciable bias. The false-negative rate will not vary depending on {*z*_*i*_, *z*_*j*_}, but the false-positive rate very likely will.

Consequently, if the target range for 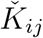 is chosen so that the *total* expected false-positive kin-pairs is no more than one or two in absolute terms, then there is not much scope for bias, regardless of covariates. The consequence of a very low overall false-positive rate will be a false-negative rate that is much higher, but the modest cost to variance (because the number of identified kin-pairs is reduced) is much preferable to a risk of substantial bias. Thus, the prescription is to first choose a target range for 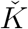 to limit the expected false-positives quite strictly, and then to estimate the ensuing false-negative rate. Then the CKMR model is implemented based on *observable* comparison-outcomes in the approximate form

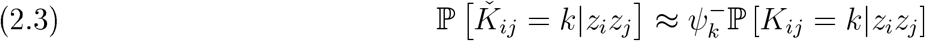

which entirely ignores false-positives. Of course, that will incur some bias but, with a strict enough limit on expected false positives, the bias can be neglected in the context of a likely-much-larger CV.

The above discussion blurs over another type of kinship uncertainty: the genetic similarity between HSPs and other 2KPs such as GGPs, which cannot be distinguished without genomic or pedigree information (Thompson, 2000; Egeland and Sheehan, 2008). CKMR probability formulae for HSPs and GGPs are quite different, and depend strongly on covariates: for example, with juvenile (immature) samples collected lethally, GGPs are impossible. The third type of non-inbred 2KP, FTP, is often guaranteed rare on demographic grounds, in species that breed repeatedly with different partners. When the covariates do not make it obvious whether a pair is GGP or HSP, the two modeling choices are either to entirely omit certain comparisons based on their covariates (because there is too much risk of multiple 2KPs and too little information gain), or to formulate the probabilities separately for HSP and GGP and add them together in the model (e.g. Thomson et al., 2020, for school shark). For the most part, the routines in `kinference` are agnostic about the target type of 2KP, leaving subsequent usage up to the modeler. However, the false-negative/false-positive calculations in `autopick_threshold` (section 4.2.5) assume that most of the kin-pairs considered will be HSPs and HTPs (rather than, say, GGPs and any other 3KPs), since these have been the most common in applications. The impact of linkage would be slightly different for other 2KPs and 3KPs; see Hill and Weir (2011) Table 2, for example.

This context should also clarify what is *not* desirable when kin-finding for CKMR, and thus not attempted by `kinference`. First, there is no attempt at probabilistic assignment, e.g. no statements of the form “this pair is 99.8% likely to be a POP” . While that is exactly the desired outcome in some kinship contexts, the problem for CKMR is that assignment requires some prior on kinship; all that can be ascertained purely from a pair’s genotypes is the likelihood of those genotypes given different kinships, so an additional prior probability would be required to calculate posterior assignment probabilities via Bayes’ theorem. However, the appropriate prior would depend entirely on the unknown demographic factors that are the desiderata of CKMR. In a way, the job of a CKMR model is precisely to *estimate* the prior probability of kinship, via explicit demographic models and estimable parameters; this would be undermined by introducing a “stealth prior” on the identification of kin-pairs. Of course, it may be desirable to place explicit priors on demographic parameters during the estimation phase of CKMR, as in any statistical analysis, but not during data setup and kin-finding.

Second, there is no single prior probability of kinship, because it will vary heavily depending on individual covariates. Although some kin-finding software (e.g. `sequoia`: Huisman, 2017) does try to formulate semi-automatic priors based on covariates and user inputs, the CKMR framework affords a more flexible and transparent way (albeit requiring user expertise) to handle the demographic aspects of kinship. We return to these points at the start of the Discussion.

## 3. Requirements and assumptions of kinference

The overall requirements for CKMR genotyping — at least within the framework of Bravington et al. (2016b) — are that it should be reliable (few errors), with enough loci to mostly distinguish 2KPs from weaker kin (though perfection is not required), but still cheap (because sample sizes can be very large). With this in mind, `kinference` handles diploid biallelic SNP genotypes for thousands of loci (but not hundreds-of-thousands). The method of genotyping does not matter, as long as other criteria are met; successful results have been obtained from targeted sequencing as well as from microarrays. Genotyping errors are tolerated, and the error rate (including allelic dropout) does not need to be known precisely, but should be low. Therefore, low-coverage genotyping is unlikely to be suitable. `kinference` does not currently use genomic information (i.e. from a high-quality de novo genome assembly) which, at least until recently, has not been affordable for CKMR species.

The data inputs comprise already-called genotypes, locus-specific information, and sample covariates (not used directly, but crucial for subsequent modeling). `kinference` expects all this in the form of a `snpgeno` object. This is a lightweight class provided by the accompanying R package `gbasics`, and `snpgeno` objects can be created directly from SNPGDS (package `SNPrelate`, https://doi.org/doi:10.18129/B9.bioc.SNPRelate) and VCF files. As the object passes through the various functions of `kinference` (allele frequency estimation, etc.), its contents are augmented.

The assumptions required for CKMR also permit some simplifying assumptions for `kinference` compared with more general kinship software, many of which can be checked with diagnostic tools in the package. Given the implicit assumption of demographic panmixia within samples, genotypes are assumed to follow Hardy-Weinberg Equilibrium (HWE). Because only a few thousand SNPs are required, the loci will not be spaced closely on average, so the impact of Linkage Disequilibrium is ignored (section 4.2.5). However, every chromosome will still contain many SNPs, so the impact of physical linkage across one or two meioses really is important and must be accommodated somehow, as described later. Genuine null alleles (repeatable and heritable; see section 3.1) are allowed for, since they have been unavoidable in some applications. However, uncalled/unknown/missing genotypes are not permitted in most `kinference` functions; every sample and locus must have a called genotype, even if some calls are wrong. The rationale is that the theoretical null/reference distributions^1^ of various statistics, which are important for diagnostics and for estimating the false-negative rate, can be calculated much more reliably and easily if every sample genotype has the same statistical properties. Sometimes, failure to call a genotype at a locus is simply a consequence of “timid” software, resolvable with more aggressive settings; if not, random imputation of a few missing genotypes should not cause major problems, since low levels of error are tolerated. Most statistics in `kinference` are likelihood-based, because that always gives the best statistical power assuming its assumptions are met (e.g. that genotypes follow HWE). However, exclusion-based statistics (modified to allow for null alleles) are also used in a few places, for simplicity and robustness.

Outputs of `kinference` are again driven by CKMR requirements. In the absence of pedigrees (which are undesirable in CKMR, as the next section explains) or genomic information, the reliably-detectable kinships are restricted to duplicates (DUPs), POPs, FSPs, and 2KPs. However, 2KPs will overlap statistically with 3KPs and perhaps weaker kin. Thus, a key part of `kinference` is the reliable estimation of false-positive and false-negative rates for 2KPs around some proposed classification threshold, making allowance for physical linkage even in the absence of a genome. Under the assumption that the genotyping data is good enough to find 2KPs, the identification of 1KPs (and resolution to POP or FSP) should be so clear that the chance of error can be neglected.

All comparisons in `kinference` are strictly pairwise, with just two alternatives of which one is usually unrelated pair (UP). This means it can take several rounds of comparisons on the same samples, with different target kinships, to identify the exact kinship of those pairs that are clearly close relatives. `kinference` does not try to automate such sequential comparisons, so by design this is a manual process. Different strategies can be employed, depending on demographics and what is actually needed in the CKMR model; the vignette has examples. Further, there is deliberately no attempt to link groups of kin together beyond the pairwise level (i.e., no pedigree reconstruction), nor to distinguish between types of 2KP (mainly HSPs and GGPs) using covariates. Although the latter is often possible, it is a case-specific issue which `kinference` again leaves to the analyst. The next subsection explains why these deliberate limitations are important for CKMR.

The ultimate output of `kinference` is partly graphical: one histogram per target kinship, where close-kin of various types should be visible as bumps at predictable locations, and sometimes with predictable shapes. If the bumps line up, then everything has worked properly, and the outcomes are the sets of kin-pairs; in the case of 2KPs, there are also estimates of false-negative probability, the false-positive probability being explicitly under user control. However, there are many reasons why the bumps might not line up, at least to begin with. So far, we have always been able to trace problems back to inadequate QC: of samples, loci, and/or the settings and interpretation of the genotyping process. While `kinference` is certainly not meant to be a full QC package for SNPs— other software, for example `radiator` (https://thierrygosselin.github.io/radiator/), has a more comprehensive scope —`kinference` does include a set of QC tools addressing issues of particular importance to CKMR (section 5). For datasets with basically well-called genotypes but perhaps other QC problems, we can successfully run entire kin-finding pipelines for CKMR using just the tools in `kinference`. Several of the QC statistics in `kinference` have null/reference distributions that can be accurately and quickly calculated by saddlepoint approximation (SPA; Appendix A).

### 3.1. Null alleles

Some genotyping methods, when applied to some species, produce a substantial proportion of null alleles. “Null allele” here means a genetic variant that fails to produce a detectable result at its locus. This lack of detection is nevertheless heritable and repeatable, e.g. if the same animal was biopsied and genotyped again. This specifically excludes allelic dropout due to non-heritable factors such as low DNA quality/quantity, PCR inhibitors, incomplete enzyme digestion, or other technical artefacts. Null alleles are important in kin-finding, because they could be misinterpreted as apparent Mendelian exclusions (Blouin, 2003). Suppose the possible alleles at a locus are A, B, and O where O is null; when a AO parent passes on its null to an o spring who inherits a B allele from the other parent, then the parent looks like an AA homozygote but the o spring looks like a BB homozygote.

Most discussion of nulls and kinship seems to focus on microsatellites (e.g. Wagner et al., 2006), and indeed the original microsatellite CKMR study of SBT found clear evidence of nulls among otherwise-obvious POPs at several loci (Bravington et al., 2016a). Nulls are less discussed with SNPs, presumably because they are not common with microarrays, and the few loci with significant null issues (e.g. from triallelic SNPs) can simply be filtered out (as suggested in e.g. Huisman, 2017, Supplement S1.1). However, CSIRO’s SNP-based CKMR projects have used sequencing instead, for reasons of cost and convenience, and we have found that nulls can be quite common in some species. For SBT, replicate genotypes and obvious parent-o spring pairs make it clear that SNP nulls are frequent (Davies et al., 2020b). In fact, given the rather small genome (0.8 Gb), large effective population size, and high proportion of polymorphic sites in SBT, it was not possible to select enough null-free loci with complexity-reduction methods akin to ddRAD for reliable kinship detection at affordable unit cost, so nulls were an unavoidable reality. In other projects, the proportion of loci with nulls has varied greatly across species; although common for other tuna, nulls have been much scarcer for rare sharks with large genomes. The decision about whether to allow for nulls in any specific project should rest on an understanding of the genotyping method in use, as well on data analysis (e.g. section 5.2). Advances in genotyping may make nulls less of an issue in future projects, but the ddRAD legacy means that allowance for null alleles is a requirement for `kinference`.

Even if nulls are likely present, not all loci will have them. While it is preferable to choose null-free loci where possible for the sake of simplicity, nulls are not necessarily a problem for kin-finding, as long as the algorithm is null-aware. For datasets where genuine nulls are plausible, the first task is to estimate the frequency of nulls as well as minor alleles (section B). Thereafter, `kinference` offers three options for dealing with nulls, which can be applied at a locus-specific level. If the null frequency for a locus is zero (estimated, or by decree), then all three options are the same.

#### 4-way

Each genotype is assigned to one of the four classes AB (heterozygote), AAO (homozygote or single-null), BBO (similar), or OO (double null). Most functions assume there are no genotyping errors across categories. This would be the usual approach if the null frequency is above a few percent.

#### 3-way

Like 4-way, except that the two categories BBO and OO are merged into BBOO . This is advisable when the estimated null frequency is below a few per cent, because it avoids over-weighting kin-pairs where both animals happen to be called OO through genotyping errors; if the double-nulls are taken at face value, this greatly in ates the odds on kinship. If nulls are rare anyway at a locus, they add little statistical power on average. Therefore, it is safer to merge the (by definition) small number of double-null calls with another category; the minimum loss of statistical power comes from merging the rarest categories, hence the choice of BBOO .

#### 6-way

This is only applicable when coverage is deliberately increased to the point where single-nulls have appreciably lower counts than homozygotes. That allows the two to be distinguished probabilistically, albeit with non-ignorable error rate (estimated in advance, and then allowed for during kin-finding). Here, nulls are functioning as bona de third alleles. Although best avoided in new projects (where more loci at lower depth for the same cost is preferred), 6-way genotyping is still used at CSIRO for SBT. Appendix F has more details.

## 4. Kin-Pair Statistics

### 4.1. Overview

All kin-pair functions in `kinference` accept two subsets of samples as arguments (possibly the same subset twice), and calculate a single statistic for each pairwise comparison. Each function calculates a different statistic that considers two-and-only-two candidate kinships, and measures their relative compatibility with each pair of genotypes, as described next.

The three basic routines, `find_duplicates`, `find_POPs`, and `find_HSPs`, have obvious purposes, except that `find_HSPs` is actually aimed at all 2KPs, and technically `find_duplicates` could also detect monozygotic siblings (identical twins, etc.), although CKMR is unlikely to be useful if those are common. Duplicate-finding is normally just a QC step, so that duplicates can be removed before further work, but it can be useful in its own right for self-recapture in live-release applications. For finding 2KPs, and in one version of the POP-finder, the statistic is a “PLOD” (Pseudo-Log-Odds, or pseudo-log-likelihood-ratio; details below) where the other candidate kinship is UP, i.e. effectively unrelated. The duplicate-finder, and the other version of the POP-finder, use more ad hoc statistics that are arguably simpler.

There is no routine specifically aimed at FSPs vs UPs, since FSPs are not usually a primary target in CKMR. In any case, unless the set of comparisons have been pre-filtered, both `find_POPs` and `find_HSPs` will usually return any FSPs as well as their main target kin. Therefore, after finding candidate POPs and/or 2KPs, the routines `split_FSPs_from_POPs` and `split_FSPs_from_HSPs` may be useful; these calculate PLODs between the two named kinships, whereas the `find_…` routines implicitly have UP as one kinship.

Once duplicates, POPs, and FSPs have been identified and removed, the final step for identifying HSPs is to deal with uncertainty in the candidate 2KPs found using `find_HSPs`, where there will often be overlap in the PLOD distributions between true 2KPs and weaker kin (3KPs and/or 4KPs). The function for this is `autopick_threshold` which, given a user-supplied tolerance for false-positives, will propose an acceptance threshold for almost-sure 2KPs (i.e. a threshold for the PLODs from `find_HSPs`) and will estimate the associated false-negative rate.

All these routines produce graphical output, mainly histograms. In the `find…` routines, to avoid the problem of storing enormous numbers (easily ∼ 10^9^, in a large project) of uninteresting statistics from every UP or almost-UP, the statistic for a pair is only retained individually if it falls above (or below) some “interesting” threshold set by the user, who can also set a cap on the maximum number of retained pairs to avoid memory over ow. The statistics for “uninteresting” pairs are aggregated into bins, as shown in later figures, since they are still important diagnostically.

The histograms from the 2KP-finder `find_HSPs` are particularly useful, since they can be used as a diagnostic to con rm that everything is working as it should— which is quite often not the case in the early stages of kinference, necessitating a return to the QC process of the next section. To exemplify the description below, Figure 4.1 shows the log-scale histogram of PLOD_HSP,UP_ for 4800 samples of school shark (*Galeorhinus galeus*) from southern Australia (Thomson et al., 2025).

**Figure 4.1.**
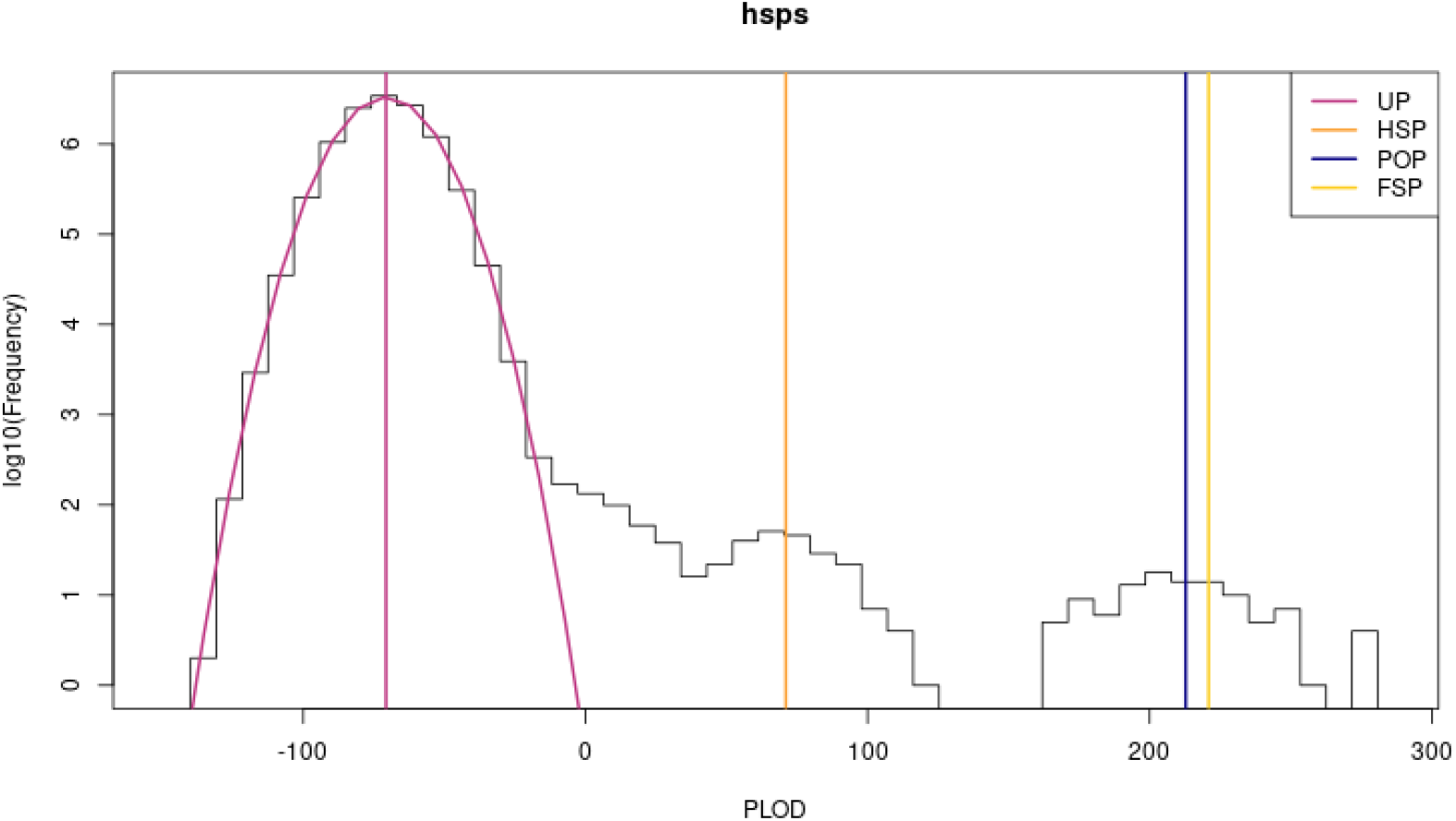
HSP/UP PLODs for school shark (as per Thomson et al., 2025), with expected values for different kinships, and entire predicted distribution for UPs.

The salient points are:

- the Y-axis is log-scale, because otherwise the vast number of UPs will compress the kin-pairs of interest (towards the right) to invisibility. (The unlogged version is also available, and is essential for zooming in on the interesting regions, roughly PLOD>0.)
- The purple lines show the predicted mean and distribution for UPs, based on allele frequencies (section 5.1). These coincide very well with observation— an essential QC diagnostic, since failure here would indicate that the assumptions of `kinference` are not being met, so that no other results could be trusted.
- The other vertical lines show expected values of the PLOD for different true kinships— not just 2KP or UP, even though those are the two candidates used in the calculating this PLOD. Only the means can be predicted, not the shapes (i.e. not variances nor heights). The POP and FSP lines are very close; this particular statistic is not meant for discriminating those two kinships.
- There are obvious bumps on the right, and they are centred around the predicted means (though in many situations, the POP/FSP or 2KP bump might be missing).
- There is also a “shoulder” between the purple UP bump and the 2KP bump, which will contain a mix of 2KPs and weaker kin.
- The expected value for UPs is close to the negative of the expected value for 2KPs— this is convenient and turns out to be generally true, although the mathematical reason is not obvious.

There is great flexibility in the sequence of kin-finding operations, which is deliberately left up to the user. There is no “just find all the close-kin pairs and report their kinship” function, because the appropriate sequence of applying the two-outcome comparisons will depend on the CKMR application; see Discussion.

The final routine directly related to kin-finding is `kin_power`, which predicts how well a proposed set of loci might work for finding 2KPs, in terms of means and variances of PLOD distributions. It is also used behind-the-scenes by many of the kin-finding routines to pre-compute useful quantities.

The following sections explain the computations behind all these functions; details of syntax are in the package documentation.

### 4.2. PLODs

For two candidate kinships *K* and *K*′, and two samples *i* and *j* with genotypes *g*_𝓁*i*_ and *g*_𝓁*j*_ at locus 𝓁, the PLOD is de ned as

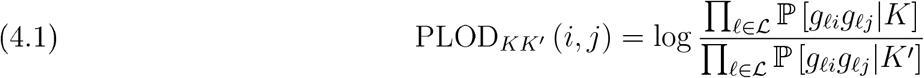

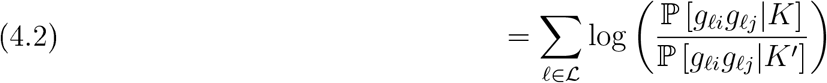

In most contexts it is clear what *K* and *K*′ should be, so we often refer just to “the PLOD”, most often for *K* = 2KP, *K*′ = UP in `find_HSPs`. The reason for “pseudo” is that the numerator and denominator in the first line are in general *not* true likelihoods, because linkage causes correlation in identical-by-descent, or “ibd”, status between neighbouring loci. However, the PLOD is still an effective measure of relative probability and it is hard to see what else could be better, in the absence of genomic information. Leaving aside the “pseudo”, the formal justification for “log-odds” is that, if *K* and *K*′ are the only two possibilities, and if they are assigned equal prior probabilities (admittedly a demographically ludicrous notion), then the posterior log-odds of *K* given the genotypes would indeed be equation (4.2). Our informal justification for preferring “PLOD” to e.g. “PLLR” (Pseudo-Log-Likelihood-Ratio) is that PLOD is more memorable and easier to say.

Following e.g. Thompson (2000), we can equate the actual kinship *K* of two animals^2^ with the three gene-identity probabilities (*κ*_2_, *κ*_1_, *κ*_0_), where *κ*_*n*_ : *n* ∈ {0, 1, 2} is the probability that *n* copies are coinherited (i.e., ibd)^3^. Apart from FSPs and DUPs, at most one copy can be coinherited at any locus of a non-inbred close-kin pair (i.e. a single-lineage pair), so mostly we have *κ*_2_ = 0. Ignoring genotyping errors and mutation for now, the probability of observing genotypes *g*_𝓁*i*_ and *g*_𝓁*j*_ at locus *𝓁* from two samples *i* and *j*, with any single-lineage kinship “XP” whose gene-identity probabilities are (0, *κ*_1_, 1 − *κ*_1_), is then

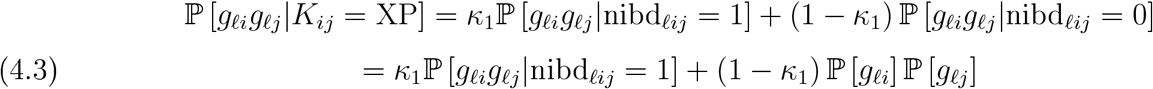

where “nibd” is the number of coinherited alleles, and ℙ [*g*_𝓁_] is the population-level probability of a genotype, computed directly from the locus’ allele frequencies assuming HWE. The term ℙ [*g*_𝓁*i*_*g*_𝓁*j*_|nibd_𝓁*ij*_ = 1] is easily computed by summing over all possible values 𝒜_𝓁_ of the coinherited allele *a*_𝓁_, as follows:

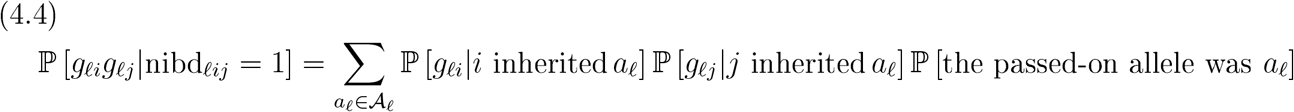

with the final term just being the frequency of allele *a*_𝓁_.

There is a natural extension to FSPs, with 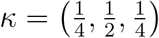, although the expressions become more tedious.

#### 4.2.1. Genotyping errors and mutations

Clearly, if *g*_𝓁*i*_ and *g*_𝓁*j*_ do not share an allele, then ℙ [*g*_𝓁*i*_*g*_𝓁*j*_|*κ* = 1] = 0; this is Mendelian Exclusion. However, a genotyping error (or, more rarely, a mutation) could lead to the *called* genotypes failing to share any alleles. Errors, provided that they are rare, make little difference for 2KPs, since the range of values of log ℙ [*g*_𝓁*i*_*g*_𝓁*j*_|*K* = 2KP] is limited so that no one locus can affect the overall result much, and only a small proportion of loci will be affected. For POPs, however, log ℙ [*g*_𝓁*i*_*g*_𝓁*j*_|*K* = POP] takes the value −∞ if there is no shared allele, so a single error could ruin an unadjusted PLOD_POP,*K*′_ . To make it robust, we can simply rede ne the gene-identity probabilities for POPs as (0, 1 − gerr, gerr) where “gerr” is a probability of pairwise error, constant across loci and samples. The assumption is that, if a genotyping error occurs, the called genotype is drawn at random from the population of genotypes at that locus; this is the “random genotype replacement” model of CERVUS 3.0 (Kalinowski et al., 2007). While this is unlikely to be an accurate model of SNP genotyping error, it does not have to be, for the same reason that errors do not matter much with 2KPs; the important thing is that it robustifies the calculation of PLOD_POP,*K*′_ while avoiding the tedium of specifying essentially-irrelevant details in a more accurate error model.

The value of “gerr” must be set by the user, based ideally on replicate samples or replicate genotypes (which need only comprise a small proportion of the total sample size, at least in a big study). The precise value should not matter much. In fact, it is a non-trivial exercise to estimate “gerr” accurately, because the random-genotype-replacement model actually implies that errors will often be invisible; the random replacement might well be the true genotype! However, a reasonable estimate might be the overall proportion, across replicates and loci that pass QC, of pairwise replicate comparisons that mismatch. Alternatively, an estimate might be based on published error rates for the genotyping technique in question. We deliberately avoid recommending a default “gerr”, because in our view the decision should be based on the user’s case-specific understanding of their genotyping approach, and also because we want to encourage the collection of replicate data. However, any vaguely-reasonable value for “gerr” should lead to essentially identical POP-finding results, because POPs are highly distinctive when there are enough loci to identify 2KPs.

#### 4.2.2. Null alleles

If null alleles are possible, then equations (4.3) and (4.4) need first to be calculated for all three possible alleles and six possible genotypes per sample, and then aggregated into observable/ distinguishable genotypes AB, AAO, BBO, or OO. This is straightforward, since the possibilities are mutually exclusive. For example:

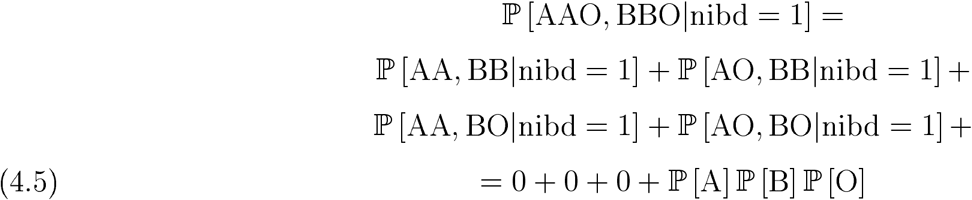

The extension to 3-way genotyping, where BBOO=∪{BBO, OO}, is immediate.

#### 4.2.3. Expectation of PLODs for different kinships

Here we use “1” and “2” instead of *i* and *j* for readability. For any *K*, the possible values of ℙ [*g*_𝓁1_, *g*_𝓁2_|*K*]can be pre-computed as a 3D array over (locus, g1, g2) since biallelic SNPs (unlike microsatellites) de ne the same set of possible genotypes G at every locus. Given two of these arrays, for kinships *K* and *K*′, the expected value of PLOD_*KK*′_can be calculated for any third true kinship *K*″, which does not have to be either *K* or *K*′:

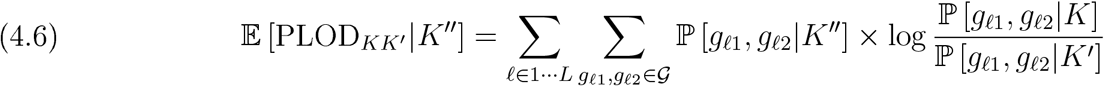

For single-lineage kinship where *κ*_2_ = 0 and 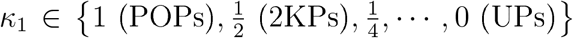, it is always the case that ℙ [*g*_𝓁1_*g*_𝓁2_|*κ*_1_] = *κ*_1_ ℙ [*g*_𝓁1_*g*_𝓁2_|POP] + (1 − *κ*_1_) ℙ [*g*_𝓁1_*g*_𝓁2_|UP], so it follows that E [PLOD|*κ*_1_] = *κ*_1_𝔼 [PLOD|POP] + (1 − *κ*_1_) 𝔼 [PLOD|UP]. Thus, for example, the expected PLOD for 3KPs is exactly midway between the expected PLODs for UPs and HSPs. In all applications that we have seen, it has turned out that 𝔼 [PLOD|2KP] ≈ −𝔼 [PLOD|UP], so that 𝔼 [PLOD|3KP] ≈ 0; in other words, we know in advance where the mostly likely source of contamination from weaker kin will be centred, which is helpful for diagnostics. The explanation seems to depend on some fortuituous near-cancellations of terms in approximate expansions of HWE genotype probabilities; we omit further details, not least because the result can always be checked directly in any application.

#### 4.2.4. Distribution of PLODs for true Ups

For *K*″ = UP, it is straightforward to calculate variance by the same route, because then the loci are statistically independent: there is no correlation of coinheritance in nearby loci, because there is no coinheritance. In fact, it is quite simple to calculate not just the variance but the entire distribution of PLOD_*KK*′_ |*K*″ = UP using SPA, because the PLOD is a sum of independent random variables each with a known discrete distribution, i.e. taking only a finite number of possible values (Appendix A). SPAs handle non-Normality well and are particularly effective at approximating far-tail probabilities without needing simulation (e.g. Butler, 2007). For the particular case of PLODs, it turns out that a Normal approximation works almost as well as the SPA, presumably because the PLOD is a sum of very many not-very-dissimilar light-tailed distributions where the Central Limit Theorem is effective. We tend to recommend the SPA version just in case, since it is better in principle.

Similar variance and SPA calculations could be done for POPs and DUPs, because coinheritance is inevitable at all loci so again there is nothing to be correlated; however, we have not seen a great need for POP or DUP variances to date. Unfortunately, SPAs will not work for other kinships, in particular 2KPs where it would be very useful, because linkage— discussed next— destroys independence between the loci. Independence is irrelevant to means, but not to variances. That is why Figure 4.1 shows the entire predicted distribution only for UPs, but just the means for the other kinships.

#### 4.2.5. Linkage

“Linkage” (e.g. Thompson, 2000, chapter 4) will be well-known to geneticists, but not necessarily to statisticians. It is the phenomenon whereby an o spring tends to inherit DNA from each parent in long chunks, each of which comes entirely from one version, or entirely from the other version, of the chromosome in that parent. Such a chunk can comprise as much as the entire chromosome, but is often shorter because of DNA recombination. During meiosis, i.e. the cell division which leads to the formation of a gamete (egg or sperm) that contains just a single version of each chromosome, the source version of its DNA can alternate ( “crossover” ) between the two versions in the parent, zero or more times across the length of the chromosome. Aside from making kin-finding more interesting, this recombination process ensures that the resulting gametes are genetically unique. It is the cornerstone of genetic diversity.

For siblings and their descendents, any chunk of DNA that is coinherited from the same version of the chromosome in their shared parent (i.e., that is ibd) will generally contain several kinship markers, since there are far more markers than chromosomes, and crossovers are only moderately common. Thus, if two half-siblings coinherit from their shared parent at the 𝓁^th^ locus along a chromosome, then they are also likely to coinherit from the same parent at the (𝓁 + 1)^th^. The probability of coinheritance between any two loci decreases as their spacing increases, and loci on separate chromosomes are inherited independently. Linkage induces substantial variability in the amount of coinherited DNA amongst close-kin pairs of the same kinship (Manichaikul et al., 2010), which contributes to the variance of PLODs for all types of kin except DUPs, POPs, and UPs. Linkage is why the 2KP bump in Figure 4.1 is relatively wider than the UP bump.

Note that linkage per se is rather different from “Linkage Disequilibrium” (LD), whereby genotypes within one individual are correlated at nearby loci, even though the two have the same underlying cause (which is that crossovers are not extremely common). In general, LD would only become significant to PLODs if many of the markers used are in close proximity along chromosomes, something which can easily be avoided when selecting the few thousand SNPs needed for finding 2KPs. Thus it is ignored by `kinference`. LD would induce correlations between LODs at different loci even among UPs, so that the PLOD variance for UPs would be higher than predicted assuming independent loci; in principle, it could be diagnosed visually from `kinference` outputs.

In this section, we refer mostly to “2KP/3KP” rather than “HSP/HTP”, but with the implicit assumption that HSPs and HTPs comprise most of the kin-pairs with PLODs in the range of interest. The reason is that the expected PLOD is determined just by the order of kinship (by definition), but the variance and higher moments are also affected by the specific type of kinship. The expected number of recombinations is higher for HSPs than for GGPs (Thompson, 2000), which reduces the effect of linkage; thus, an HSP bump would be narrower than a GGP bump for the same species. Assuming the main CKMR focus is HSPs not GGPs, it is therefore best to restrict 2KP comparisons a priori to pairs unlikely to be GGPs (based on likely birth-gap, say), so that linkage variance is estimated more reliably. Nevertheless, a modest proportion of GGPs among the HSPs seems unlikely to cause major biases in kin-finding (though it would need to be allowed for in CKMR modeling per se). And if the birth-gap among the comparisons used is low enough to largely exclude GGPs, then HTPs (whose average birth-gap equal to the generation length) will be much more common than the other single-lineage 3KP (GGGPs, or Great-Grandparent-Grandchild Pairs, whose average birth-gap is three generation-lengths).

#### 4.2.6. Threshold selection for HSPs

The impact of linkage on PLODs depends on the species (and sometimes also on the sex of the shared parent; see 4.2.8). It reflects how the genome and the available markers are organized into chromosomes, and on the rate of recombination (which in fact varies within chromosomes, but for our purposes can be approximated by an overall average rate). The traditional way to estimate that rate is by building a “linkage map” based on an extensive known pedigree across several generations, something which is not feasible for most wild animals. However, the practical impact of linkage on HSPs can be readily inferred by comparing the observed bump width, i.e. variance, of HSPs to a zero-linkage prediction obtained by summing the individual per-locus variances. Since the distribution of PLODs for HSPs tends to be quite Gaussian, the empirical variance estimate can then be used to predict the false-negative proportion of HSPs below any given threshold *τ* .

One practical issue with estimating the “2KP bump width” is identifying a set of clearly-2KP kin in the first place, given the overlap with 3KP and other kin. Fortunately, the upper tail of PLODs from 3KPs (or any specific kinship) decays rapidly, and unless the study is badly designed, the mean 2KP PLOD should be well above the likely range of 3KPs, which should be verified visually. Our normal policy is to use all *n* PLODs *y*_*ij*_ above the 2KP mean of *e*_2KP_ (and below a threshold to exclude first-order kin, which should be visually obvious), and to estimate the variance of a half-truncated Normal distribution from *n*^−1^ Σ (*y*_*ij*_ − *e*_2KP_)^2^. Sometimes it might make sense to use a higher cut-o (if bump separation is poor, to more stringently exclude possible 3KPs) or lower cut-off (if bump separation is good, to increase the number of 2KPs for variance estimation), and then to use the formula for the variance of a general truncated Normal.

The more challenging question is how *τ* should be chosen so as to control the number of false-positives from 3KPs (4KPs are unlikely to contribute directly for any reasonable choice of *τ*, but still have an indirect role to play in estimation, as explained later). That false-positive number depends on:

1. how many 3KPs are truly present in the sample;
2. the PLOD variance for 3KPs;
3. the PLOD mean for 3KPs.

The last can be calculated in advance, as per the last subsection. The other two must be estimated from the available data: namely, the observed PLODs in the range where 3KPs and 2KPs will overlap. That range is at the user’s discretion, but might lie roughly between 0 (the 3KP mean) and the 2KP mean.

The function `autopick_threshold` fits the PLOD distribution in that range using a Normal mixture with either two or three components, to describe 2KPs, 3KPs, and perhaps 4KPs which may intrude near PLOD=0. For 2KPs, the mean and variance can be pre-estimated as above, but for 3KPs and 4KPs only the means are known in advance. However, the observed 2KP variance gives information about the overall impact of linkage, from which it is possible to calculate bounds on the 3KP and 4KP variances, under two extreme assumptions about genomic organization:

1. there are *C* similar chromosomes, each containing a proportionate mix of all kinship loci, with no recombination;
2. there is just one huge chromosome, with kinship loci equally spaced and in random order, and a recombination rate *ρ* between loci.

Under each assumption, the unknown parameter (*C* or *ρ*) can be estimated from the empirical 2KP variance, and then used to extrapolate the variance for 3KPs and 4KPs. As noted earlier, it is also necessary to assume that the 2KPs are mostly HSPs, the 3KPs are mostly HTPs, and the 4KPs are mostly HCPs. The variance calculations are given in Appendix E, and follow similar lines to e.g. Hill and Weir, 2011. The two versions of extrapolated variances presumably span the truth. In our experience, the second assumption (one big chromosome) always gives slightly higher variances for 3KPs than the first (many small chromosomes).

To implement this, the function `autopick_threshold` first calculates the variance bounds as above, then chooses (say) ten equally-spaced values across those ranges in parallel (so that if e.g. the 5th possible value of 𝕍 [PLOD|3KP] is being used, then the 5th possible value of 𝕍 [PLOD|4KP] will also be used). For each of the ten possible variance sets, `autopick_threshold` then fits by maximum likelihood a Normal mixture model with *known* means and variances, to estimate the relative heights of the bumps for 2KPs and 3KPs, and for 4KPs if used. For each such t, *τ* is calculated so that the expected total number of HTPs with PLOD<*τ* equals the user-supplied tolerance. The overall suggestion for *τ* is then either the *τ* for the best-fitting of the ten models, or— for a particularly cautious user— perhaps the largest *τ* across any of the models.

Even though some fairly intricate statistics are involved in the computation of potential *τ* -values and associated false-negative probabilities, we emphasize that the final selection of *τ* is an “engineering tolerance” decision, not the inevitable result of a well-de ned estimation problem. Provided that the predicted number of false-positives is below whatever tolerance the user deems acceptable, then any larger value of *τ* could reasonably be chosen; building the *τ* -specific false-negative probability into the CKMR model will compensate to eliminate bias, although there is a trade-o in increased variance as *τ* is increased. One way to emphasize this harmless subjectivity is to round the proposed *τ* upwards to a convenient nearby number, making it clear that *τ* is not an estimate but more of a safety factor. If the number of definite 2KPs remaining is so small that the choice of *τ* really matters, then one might question whether there will be enough to t a useful CKMR model anyway.

It would in theory be possible (and simpler) to t the mixture model without constraints on variance, i.e. just allowing the 3KP and 4KP variances to be additional free parameters. However, in our general statistical experience, mixture models are prone to misbehaving when given too many free parameters, so it pays to constrain the fitting using as much prior information as possible.

#### 4.2.7. Example of PLOD threshold selection

We return to the school shark example of Figure 4.1. Given the likely ages and capture-dates of the samples, most 2KPs should be HSPs and most 3KPs should be HTPs, so the assumptions of `autopick_threshold` should be reasonably satisfied. Figure 4.2 shows the unlogged histogram of PLODs, zoomed in to PLODs of -10 and above. The curves show expected counts from the best-fitting Normal mixture, using only PLODs of 0 and above, with the means and variances constrained as above. The proposed threshold *τ*, just above PLOD=50, corresponds to a user-chosen tolerance of 1 expected 3KP with PLOD *> τ* .

**Figure 4.2.**
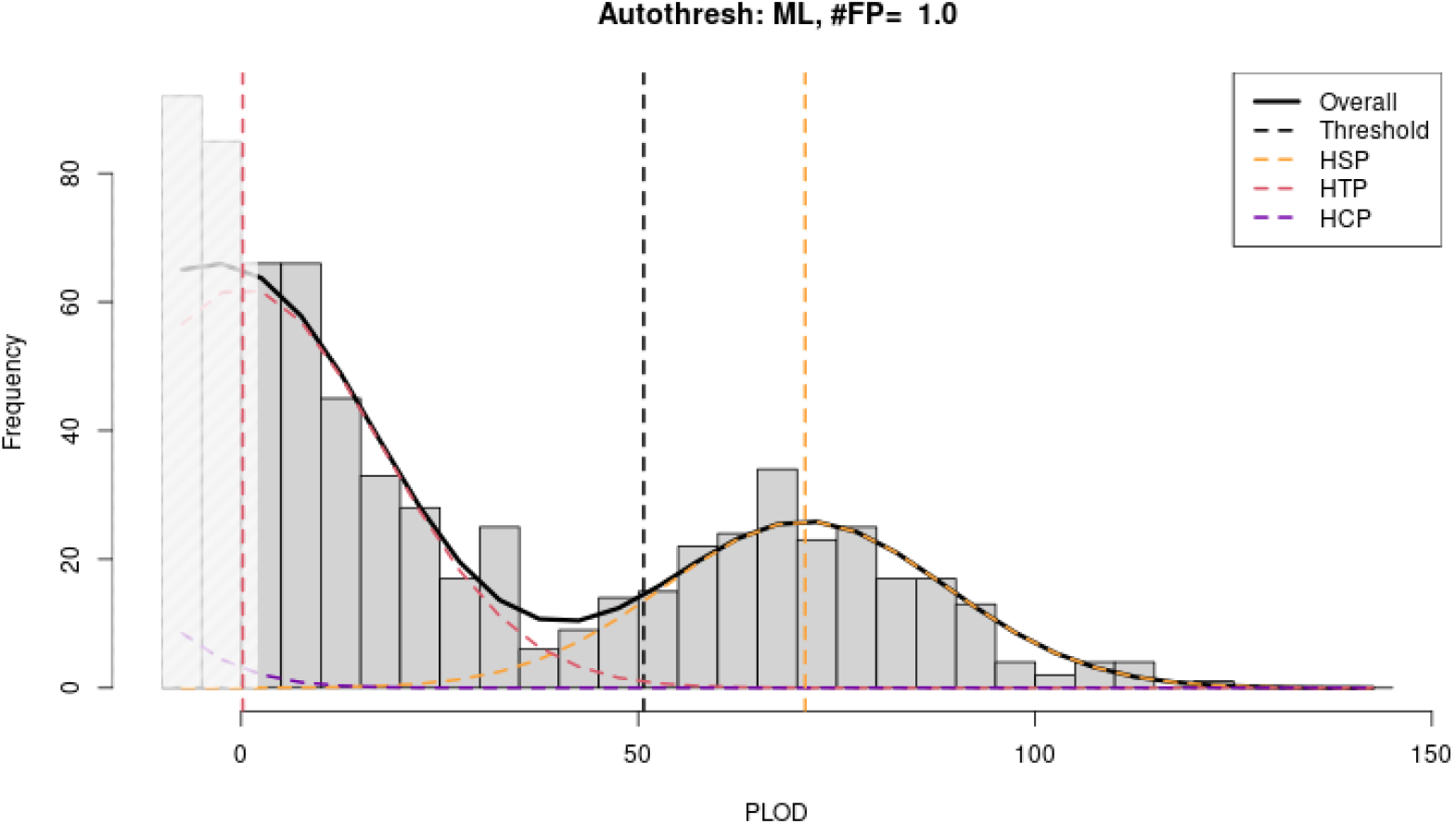
PLOD threshold selection for 2KPs in school shark, (as per Thomson et al., 2025); see text for details.

The t is good over the range of PLODs actually used. The Normal-mixture model clearly under-predicts the number of pairs with PLOD just below 0 (which were not used in fitting), but that is not surprising; those pairs are demographically likely to include large numbers of 4KPs and even 5KPs, in quantities that cannot be reliably predicted from PLODs above 0. In this case, 4KPs evidently have a very low probability of PLOD>0, so there is not much information to extrapolate their total number at lower PLODs. Overall, this particular t seems good within the range that matters, and the input tolerance for false-positives is quite conservative, so a threshold of 50 would be reasonable; given that there are about 200 presumably-2KPs above the threshold, CKMR estimates are unlikely to be biased much if 1 or 2 or 3 contaminating 3KPs do manage to sneak into the “definite 2KP” set. The estimated false-negative rate with *τ* = 50 is just 12.5%, so rather little information is being wasted in the discarded HSPs.

#### 4.2.8. Heterochiasmy

Crossover rates may di er between eggs and sperm; this is heterochiasmy. The extent varies substantially between species (Lenormand and Dutheil, 2005), and is not easy to predict. For kin-finding, heterochiasmy means that the width of the HSP bump may di er for maternal and paternal HSPs, and similarly for HTPs depending on the sex of the shared ancestor. Estimating an overall variance for all HSPs, and choosing a common cutoff, may lead to sex-specific inaccuracies. One x would be to use mtDNA to check the sex ancestry, and then to do kin-finding separately for same-mtDNA and different-mtDNA HSPs, with two distinct cutoff s and false-negative rates.

### 4.3. Power of loci for kin-finding

Although not strictly part of kin-finding per se, the PLOD calculations in `kinference` are also useful for deciding whether a proposed set of marker loci will be adequate for kin-finding, in particular for HSPs. In most CKMR projects that we have seen, the genetic work ow consisted of a ddRAD SNP discovery step, followed by marker selection based on minor allele frequency estimates as well as numerous QC parameters, for inclusion in a more cost-effective targeted assay. In our experience, a genome assembly will improve the targeted assay design process and marker conversion rate (i.e the number of markers successfully targeted, divided by the total number of markers proposed for the assay), therefore alleviating the need to build redundancy in the assay in order to compensate for the failure of some of the markers.

But how many loci should be included in that targeted assay? Using too few is catastrophic, because there are vastly more UPs than true 2KPs, and with too few loci the right-hand tail of PLODs from Ups will swamp the PLODs from true 2KPs. On the other hand, genotyping costs do scale to some extent with the number of loci, and eventually exhibit diminishing returns, whereby the ability to reliably distinguish 2KPs does not increase much by adding more loci. The limitation is intrinsic variation in the amount of coinherited DNA, driven by linkage (e.g. Manichaikul et al., 2010). These considerations can be explored numerically via the `kin_power` function, as follows.The more loci used, the greater the distance between the mean PLOD_2KP,UP_ for true UP and true HSPs. Since the variance of the UP bump in Figure 4.1 is also predictable, and the number of UPs will roughly equal the number of comparisons (only a small proportion of comparisons will yield a close-kin pair), it is not hard to check for a set of loci whether, say, the far-right hand tail of the UP distribution will reach anywhere close to the 2KP mean. Larger sample sizes might suggest that more loci are needed, because the UP distribution will spread further; in fact, though, that effect is surprisingly small because the UP distribution is well-approximated by a Normal distribution, and thus is very light-tailed.

However, despite the enormous numbers of UPs in many studies (e.g. ∼ 10^8^ for SBT), experience has shown that the more serious PLOD-overlap problem in kin-finding is usually between 3KPs and 2KPs, because their means are much closer together. Here the impact of linkage really matters, because it determines how much the 2KP and 3KP PLOD bumps will overlap. Unfortunately, that will not be known at the time when candidate loci are being chosen. The best option seems to be to use a conservative rule-of-thumb, perhaps as follows.

We have found empirically that the PLOD variance for HSPs has generally been 3–4 times higher than for UPs; although this depends on the species and will not scale closely with the number of loci, it is at least roughly useful. The HTP variance is between that of HSPs and that of UPs, because the linkage-driven variability in proportion of coinherited DNA matters less when that proportion is smaller in the first place, so perhaps 2–3 times higher than the UP variance. It will be more conservative (safer) to assume higher values. The UP mean is close to the negative of the HSP mean, and the HTP mean is therefore close to zero. The number of HSPs and HTPs will generally have similar order-of-magnitude though this can vary, depending mainly on sampling strategies. If to be safe we assume that the true number of HTPs is about twice the number of HSPs, and if we are designing the sampling in the expectation of there being 100 true HSPs, then a conservative false-positive threshold that on average includes only 1 HTP should be at a PLOD of around 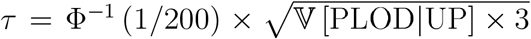, where Φ() is the cumulative distribution function of a standard Normal. The corresponding false-negative probability for HSPs would be 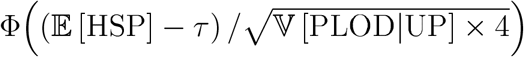. If this is unacceptably high for some proposed set of loci, then it would be wise to add more.

Ideally, similar calculations should be made in a case-specific way consistent with the CKMR model, taking account of likely demographics and sampling patterns to give some idea of the true 3KP/2KP ratio for the comparisons that will actually be used. In practice, a rough and discrete decision might be acceptable: for example, whether 2500 loci would be sufficient given the estimated distribution of minor allele frequencies, or whether 3500 would be safer. At any rate, the calculations can be explored with the function `kin_power`, which takes a real or hypothetical set of allele frequencies (minor and null) and returns expected means and variances of PLOD_UP,2KP_ for UPs, and the means for 2KPs and POPs.

### 4.4. Kinship statistics not based on likelihood

For DUPs and POPs, `kinference` offers non-likelihood-based options. `find_duplicates` simply counts the number of non-identical 4-way genotypes in each pair. Even true duplicates will tend to have a few mismatches because of genotyping errors, but usually there is an obvious gap between a spike of true duplicates at or close to 0 mismatches, and the next clump of pairs (most likely cormprising POPs and/or FSPs). If not, there may be underlying QC problems.

For finding POPs, Mendelian exclusion— checking that there is a shared allele at every locus, allowing perhaps for errors— is a traditional approach (e.g. Jones and Ardren, 2003). It is simple, easy to explain, and robust to genotyping errors: a few apparently-excluding loci will not destroy a statistic such as “number of apparently-excluding loci”, whereas an unrobustified PLOD will go infinite. Although exclusion is in principle less statistically efficient than likelihood^4^, efficiency is not really an issue for POPs when the number of SNP loci has been chosen with the much-less-detectable 2KPs in mind; POPs should basically be obvious. However, it was not always thus. The first, POP-only, version of CKMR for SBT used microsatellites, as the only proven technology in 2006. Dealing with non-rare and non-simple genotyping errors, null alleles, and unscorable loci was a major challenge that would be even worse in a likelihood setting where detailed probabilistic models would be needed, so POPs were instead determined from an exclusion-based statistic adjusted for null alleles. When the SBT project moved to SNPs, it was natural to keep going with the exclusion-based statistic. For new projects, we tend to prefer PLOD_POP,UP_, but it does no harm to try both approaches.

The major complication with exclusion for SNPs is null alleles. If ignored, then two apparent homozygotes of opposite type would constitute a clear exclusion, whereas in fact an AAO and BBO might coinherit the null allele. The null frequency varies substantially across loci, so that such “pseudo-exclusions” are quite informative (i.e. suggestive of true exclusion) if the null frequency is low, but less so if the null frequency is high. `kinference` therefore calculates a “Weighted PSeudo-Exclusion” (WPSEX) statistic, as a weighted sum across loci of the number of pseudo-exclusions:

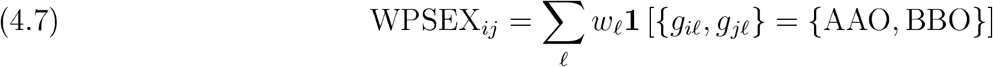

where **1** () is the indicator function (i.e. 1 if its argument is true, or 0 if not). The weights *w*_𝓁_ are chosen to minimize one type of false-positive: that an UP might have a WPSEX as low as the average WPSEX for a true POP. See Appendix C for details.

Another kind of exclusion occurs if {*g*_*i*_, *g*_*j*_} = {AB, OO}. This is not a “pseudo” exclusion since, in the absence of errors, it should never happen for POPs, although it should not be very common even for UPs because loci with high null frequencies (required to generate a lot of double-nulls) are best avoided. Nevertheless, the number of such cases— the “NABOO” statistic in kinference— does add some power to POP-finding, for example in distinguishing between POPs and FSPs.

#### 4.4.1. Example of finding POPs

Figure 4.3 compares the PLOD and WPSEX results for a subset of SBT samples (dataset `bluefin_4` in the package vignette). In the PLOD figure, it turns out that the clear bump on the right-hand side contains seven pairs, all with PLOD_POP,UP_ close to the expected value for true POPs. These are the same seven points in the lower-left of the WPSEX figure, including the lonesome dot close to (0.04,4). However, running these seven candidates through `split_FSPs_from_POPs` reveals that the lonesome dot is clearly an FSP not a POP. It has PLOD_FSP,POP_ = +43, which is close to the expected value of +47 for FSPs, whereas the other six pairs all have values of -26 or below, compared to the expected value of -37 for POPs. With that one pair excluded, both figures show a clear gap between the same set of six POPs and all other pairs. Note that the 2D plot with NABOO as well as WPSEX is helpful: none of the six true POPs have NABOO>2 (which must be from genotyping errors) amongst the 1541 loci, but the solitary FSP has NABOO=4.

**Figure 4.3.**
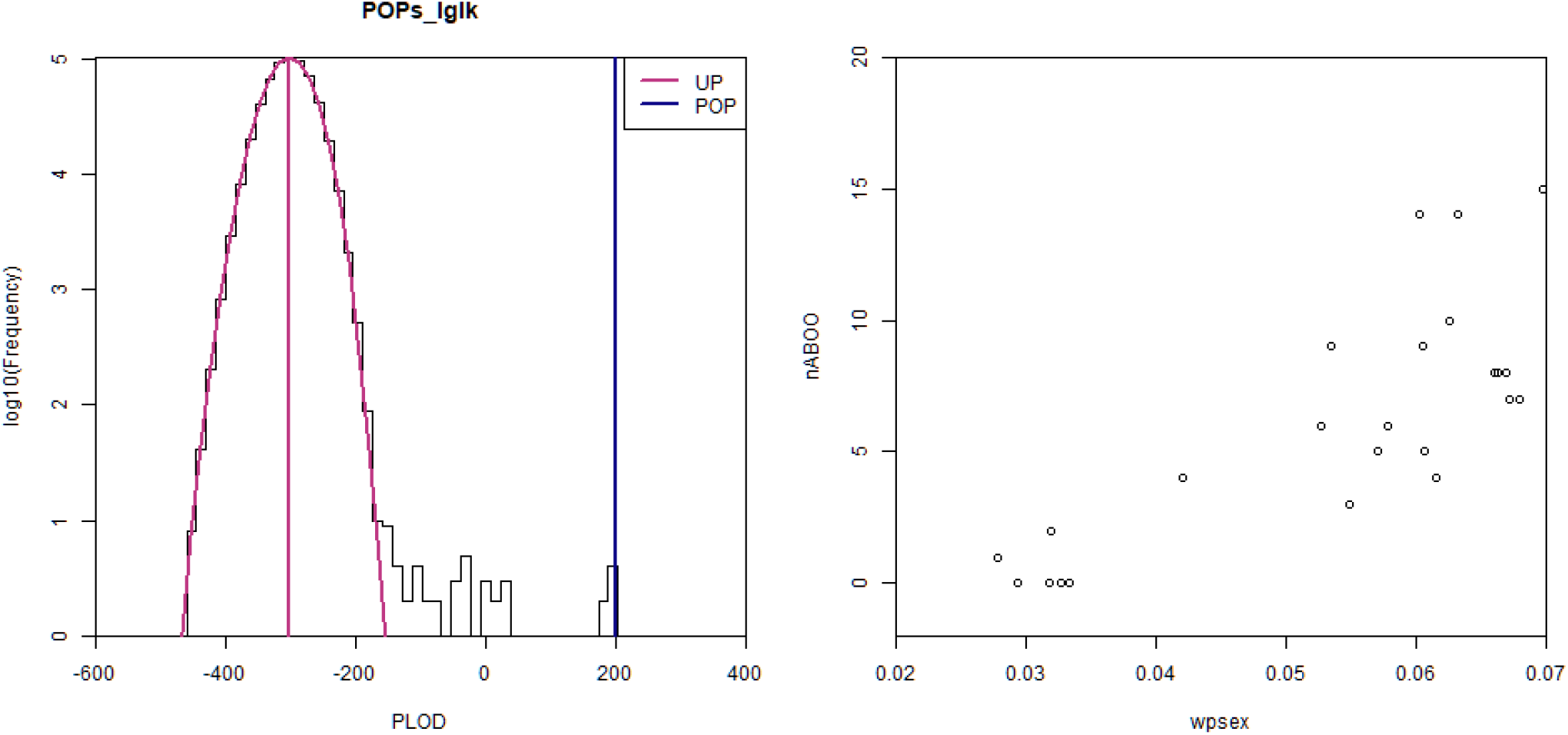
Finding POPs for a subset of SBT using PLODs (left) and WPSEX/NABOO (right). Dataset `bluefin_4` from the package vignette. The “gerr” parameter was set to 0.01 for the PLODs.

## 5. The QC cycle

Careful QC, both of loci and of samples, is crucial to successful kin-finding. The ultimate test is the log-scale PLOD histogram from `find_HSPs`, as in Figure 4.1; if this looks the way it should, and if there are enough true kin-pairs visible to check consistency, then the intervening steps have self-evidently worked. However, there are quite a few of those intervening steps, and each one can go wrong and needs to be checked by a human. Because a few bad samples can make innocent loci look guilty, and vice versa, QC for kinference is fundamentally iterative, provisionally removing and re-introducing suspect samples and loci until— hopefully— a satisfactory outcome is achieved. If not, or if too few samples and/or loci remain, then the problem may lie among the many earlier steps of data preparation, starting from in-the-field data collection through the various stages of preparation and genotyping. Such problems are sometimes easy to find and fix (e.g., if the genotype-calling software is making too many “missing” calls when it is just not sure), but in general are beyond our scope here.

Our QC philosophy for `kinference` is deliberately subjective and visual, at least until the very last steps. The QC functions in `kinference` provide statistical evidence on which provisional decisions can be made about what to include or exclude, but there are no default parameters, and it is up to the user to inspect the results and make their own decisions. The general idea is to shrink the data to a core set of clearly-good loci and clearly-good samples, removing duplicates along the way, and then to confirm that `find_HSPs` (or the PLOD version of `find_P0Ps`, if POPs are the only goal) is working properly; if not, something more fundamental has gone wrong. Once `find_HSPs` is working well on a perhaps-overly-restrictive subset of data, the next step is to try re-admitting some samples that might have been unfairly excluded because of bad loci, and vice versa. The whole process may need to repeat a couple of times, and allele frequencies should be re-estimated whenever the working set of samples is changed. Once the dataset is stable, POPs, FSPs, and HSPs/2KPs can be found with the tools of the previous section. There should then be a check of kin-pairs for internal consistency (section 5.5), which might lead to revisiting earlier steps.

In our view, QC for kin-finding is (and will continue to be) a partly subjective process that benefits from experience. Nevertheless,while there is some scope for different analysts to be more or less generous about what to include at each stage, the iterative process of selectively re-admitting samples and loci, plus the use of estimated false-negative rates to compensate for PLOD threshold choice, helps make the final outcome of CKMR largely robust to subjective choices made when running `kinference`. Small differences in chosen thresholds and included samples can slightly affect the number of kin-pairs found and the estimated false-negative rate, but in the context of CKMR itself, where parameter CVs have never been below 10% in our experience, variations of a couple of percent in the “true” number of kin-pairs used are not important.

### 5.1. Allele frequency estimation

`kinference` includes three functions for allele frequency estimation, of which one is just for 6-way genotypes and is covered in Appendix F. Otherwise, for users confident that true nulls are not present, `est_ALF_nonulls` simply counts A- and B-alleles per locus, ignoring any OO genotypes (which, in this one instance, are assumed to be “don’t know” rather than true nulls) and treating AAO and BBO genotypes as definite homozygotes. For datasets where true nulls are a realistic possibility, the basic tool is `est_ALF_AB0_quick`, which finds MLEs of minor-allele and null-allele frequency for each locus (i.e. two free parameters). It uses a modified EM algorithm, ameliorating the latter’s notoriously sluggish convergence speed by Aitken acceleration (Appendix B). `est_ALF_AB0_quick` is very fast, converging for thousands of loci in just a few seconds, despite being coded in native R. This is mainly because it can be readily vectorized; earlier versions using direct MLE per locus ran perhaps 100 times slower.

Accurate estimates of allele frequencies seem to be most important for checking overall dataset QC (i.e., the match between observed and predicted distributions of various QC statistics), and reasonably important for kin-finding; outlier detection tends to be less sensitive, so for example it may not be worth trying to remove duplicates before using `est_ALF_AB0_quick` for the first time . If `est_ALF_AB0_quick` is used on a locus without true nulls (and also without any mistaken OO calls), then it will often estimate some small but non-zero null frequency, simply because the freedom to estimate another parameter may allow a slightly better fit to the AB, AAO, and BBO counts. Since null-frequency estimates cannot be negative, but might be positive due to such noise, this represents a systematic bias for loci without true nulls; the bias diminishes with increasing sample size, because sampling variability is reduced. No single-locus statistical test could ever distinguish between zero and very-low null frequency, but when all loci are considered together, a modest impact on QC statistics can be seen in simulated data where nulls are not “really” present but `est_ALF_AB0_quick` is used instead of `est_ALF_nonulls` (e.g., in the `dropbears` example in the vignette).

In practice, we have simply used `est_ALF_AB0_quick` (or its ancestors) on most datasets, unless we were *a priori* sure that nulls were negligible. In principle, this is not strictly optimal since, even if the overall genotyping technique is capable of generating true nulls, many loci will still be genuinely null-free. Perhaps some kind of shrinkage estimator of *π*_O_ could do better, setting 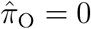 exactly unless there is strong enough evidence to the contrary for a locus, and taking all loci into account. However, we have generally had satisfactory kin-finding results from `est_ALF_AB0_quick`, once any underlying QC issues have been resolved.

### 5.2. Per-locus genotype frequencies

For each locus, the number of AB, AAO, BBO, and OO genotypes in the sample can be predicted from the allele frequency estimates (minor and null) assuming HWE, and then compared with the observed totals. The overall goodness-of-fit for the locus can be summarized by a G-statistic 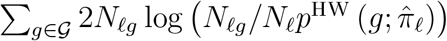 where *N*_*ℓg*_ is the number of samples with genotype *g* at locus *ℓ*, and *N*_*ℓ*_ ≜ Σ_*g*∈𝒢_ *N*_*ℓg*_ is the total sample size, and *p*^HW^ (*g*; *π*) is the probability of observed genotype *g* given allele frequencies *π*. If 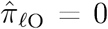 exactly, then *p*^HW^ (OO) = 0 and there cannot be any observed double-nulls, so then summation should exclude *g* = OO. We have found the G-statistic to be a little more stable than the common Pearson alternative, though either could be used. It should approximately follow a 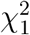 distribution since there are 4 categories, 1 fixed sum, and 2 estimated parameters. Calculations and graphics are handled by `check6and4` (so named as it was originally written for 6-way genotypes, as in Appendix F). Figure 5.1a shows an example for a subset of SBT data, including many loci that are clearly not working properly and should be (and were) discarded; it is discussed further below.

**Figure 5.1.**
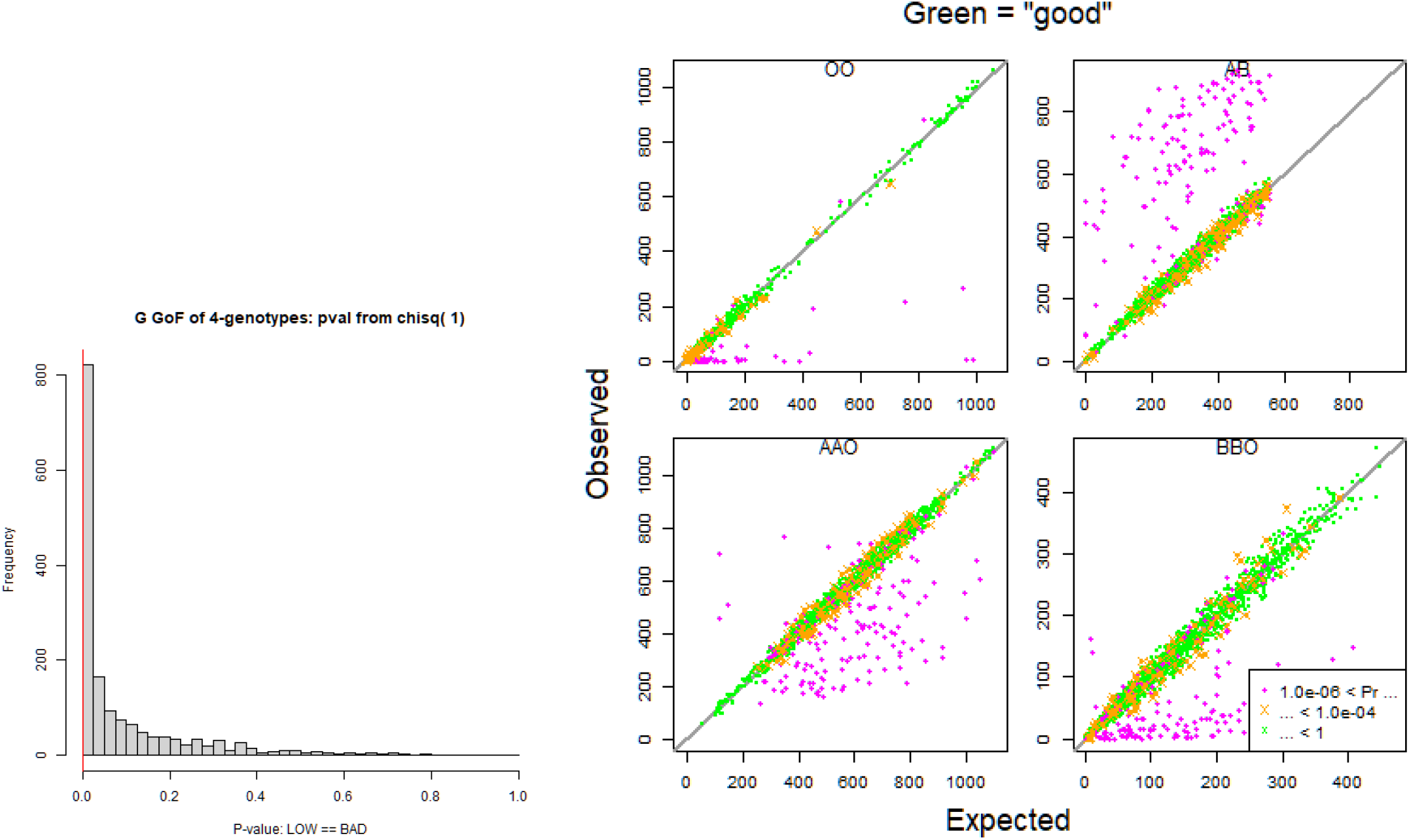
Goodness-of-fit by locus of 4-way genotypes for SBT, *before* QC. (a) Observed and predicted counts by genotype, with each locus being one dot in each panel; (a) distribution across loci of overall *p*-value. Pink is bad, green is good.

If all underlying assumptions are fully met, then the *p*-values for well-behaved loci should be uniformly distributed on [0, 1], but in practice we have found them to be concentrated towards low values. This is clear in Figure 5.1b, even ignoring the spike near 0 that comes mostly from really bad loci which will be discarded. The effect has been especially apparent in large datasets where, of course, sample size will increase the significance of *any* departure from HWE and/or from the 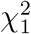 approximation, regardless of the magnitude of that departure. The issue is not so much multiple-testing (with thousands of loci, a few are bound to have to *p*-values below 0.001) but that the underlying assumptions behind the *p*-value are often *slightly* incorrect. The bigger the sample size, the worse the *p*-values will look. Nevertheless, kin-finding has often worked well despite including many loci with ostensibly extreme *p*-values, in terms of the overall PLOD distribution fitting well to theory (e.g. Figure 4.1). Thus, the *p*-values do not provide any absolute measure of “badness for kin-finding”. However, they are still useful for ranking loci and rejecting bad ones below some (subjective) cutoff.

The main diagnostic of locus adequacy is not the *p*-values per se, but the observed-vs-expected plot, which serves two purposes:

1. diagnostic of “broad-spectrum” problems, based on overall patterns;
2. indication of bad loci, as outlying points.

Broadly speaking, the plot for each genotype (AB, AAO, etc.) should be a scatter around the *y* = *x* line, sometimes with obvious outlying loci. Systematic departures from the straight line are indicative of large-scale problems. As just one example: if a substantial proportion of genotypes are simply unscored but have (mistakenly, from the point of view of `kinference`) been recorded as double-nulls, then the estimated null frequencies will be quite high, because the probability of a true double-null is the square of the null frequency; that appreciably reduces the estimated A- and B-frequencies, so that the observed AB counts will tend to sit above *y* = *x*.

Perfection is not to be expected, and some artefacts arise simply through the combination of an HWE assumption and estimated allele frequencies, especially when nulls are possible. For example, it is common for the AB scatterplot to sit somewhat above *y* = *x* at its far right-hand end. The reason is that the *expected* number of heterozygotes can never exceed half the number of samples, whereas the observed can do so simply by chance; and if the observed falls below that line, then the null-frequency estimate will tend to rise to compensate, in effect moving the point leftwards towards the line.

Regardless of overall goodness-of-fit, bad loci often stand out quite obviously. For example, paralogous loci have anomalously high numbers of apparent heterozygotes, so will sit well above the *y* = *x* line. The `threshold` parameter of `check6and4` can be used to highlight those loci with lower *p*-values (pink dots in Figure 5.1a). Our policy is to experiment with different thresholds until a visually satisfactory effect is obtained, then provisionally exclude highlighted loci. This forms part of the iterative QC cycle.

Although it is not a good idea to take literally the absolute *p*-values from `check6and4`, the scale of the *y*-axes in Figure 5.1a is meaningful. Roughly, if HWE applies then the count of any one genotype for one locus should be Binomially distributed, thus with variance somewhere between the expected value (i.e., the *x*-value) and half the expected value. Thus, *y*-values outside 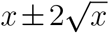 are perhaps worth paying attention to, but smaller departures are certainly insignificant.

There is an art to interpreting these plots, in terms of “how bad is too bad”. In the end, the real test is not so much in the output of `check6and4` itself, but rather in PLOD histograms like Figure 4.1. Excluding too many loci will reduce overall kin-finding power, so that more true 2KPs will be have to be sacrificed in order to safely exclude weaker kin; however, including too many bad loci can lead to the worse problem of mismatches between the observed and predicted locations of PLOD bumps, so that false-negative rates cannot be estimated reliably.

### 5.3. Per-sample log-likelihood

A useful first-pass QC statistic is to calculate the log-probability Λ^ilglk^ of each sample’s entire genotype, i.e.

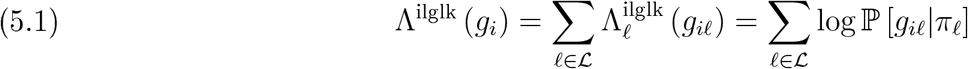

This is effective at, for example, detecting samples inadvertently included from other genetically-distinct populations, or from closely-related species. The null distribution is readily calculated, since the loci are independent by assumption. The moment-generating function (MGF) for one locus, *ℓ*, is

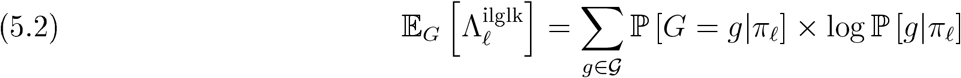

which can be used for SPA as in Appendix A.

This is handled by the function `ilglk_geno`; a typical output is shown in Figure 5.2. Outlying samples usually lie in the left-hand tail (e.g., from poor-quality DNA, or even from a closely-related species that has been mis-identified). It is common to have some discrepancy between the null distribution and the overall observed distribution. However, unless severe (a matter of judgement), we have often found that such a dataset will still work well for kin-finding, in terms of observed and expected PLOD distributions. In essence, `ilglk_geno` appears sensitive to some imperfections that do not have much impact on PLODs. It is possible that a more nuanced statistic could be devised to do better (i.e., not to show unimportant misfits) but `ilglk_geno` is only meant as a general-purpose first-pass statistic, for which we have found it useful.

**Figure 5.2.**
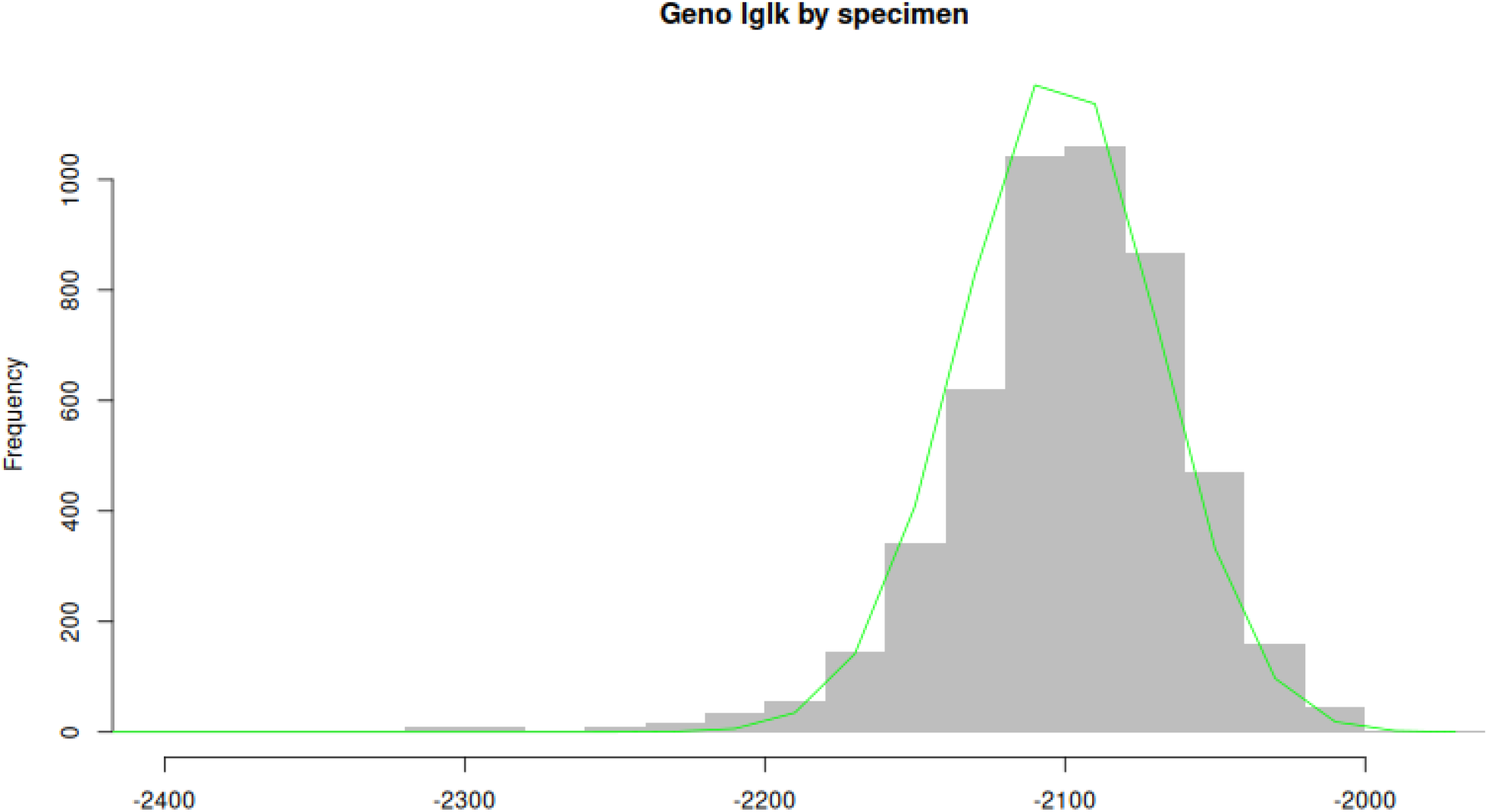
Distribution across samples of overall log-likelihood (ilglk_geno). Example for school shark, as per Thomson et al. (2025).

### 5.4. Per-sample heterozygosity checks

Abnormal levels of apparent heterozygosity in a sample are a possible indicator of contamination (because a homozygote or single-null in the source DNA may be scored as a heterozygote after contamination by another sample) and of degraded, poor-quality DNA (because one of the alleles in a source heterozygote may drop out, so that it looks like a homozygote or single-null). Similar considerations apply to apparent double-null genotypes: true double-nulls are less likely to be scored as such in contaminated samples, and single-nulls especially are likely to be scored as double-nulls in samples with dropout.

To combine all this into simple diagnostics, we first define a single-locus “hetzminoo” statistic (heterozygotes minus double-nulls), and then aggregate it across all loci for each sample. The single-locus statistic is *S*_*ℓ*_ ∈ {−1, 0, 1} ≜ **1** [*G* scored = AB] − **1** [*G* scored = OO]. Its expected value and variance, *e*_*ℓ*_ (*θ*) and *v*_*ℓ*_ (*θ*), are given by

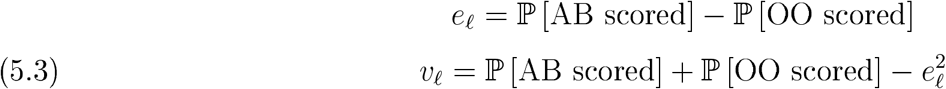

The idea is that *S*_*ℓ*_ in a contaminated sample is more likely to be +1 and less likely to be -1, and conversely in a poor-quality sample. The distribution of *S*_*ℓ*_ depends of course on the contamination/ degradation level, but also on the minor allele frequency (MALF) and null allele frequency (NALF) of locus *ℓ*. If MALF=0.99 then very few samples will be scored as heterozygotes even when contamination is high; however, if a sample does have many heterozygotes among such loci, then contamination is more strongly indicated than by the same number of heterozygotes among loci with lower MALF.

The aggregate statistic for sample *i* is a weighted sum of its single-locus *S*_*ℓ*_ values. That is, we calculate *H*_*i*_ (*w*) = Σ*w*_*ℓ*_*S*_*iℓ*_, with the weights *w*_*ℓ*_ chosen based on MALF and NALF. For each sample, the null hypothesis is that it is uncontaminated and undegraded, with all genotypes following HWE. We can choose optimal weights that should give high statistical power to a specific alternative hypothesis about “error”. Thus there are two versions of *H* (*w*), i.e. two different sets of weights, one focused on contamination (where higher values are a bad sign) and the other on DNA degradation (where lower values are a bad sign). Appendix D shows the derivation of the optimal weights, and also how the null/ reference distribution of *H* (*w*) can be obtained by SPA. Figure 5.3 shows an example of the “degraded” version of the hetzminoo statistic, before and after data cleaning.

**Figure 5.3.**
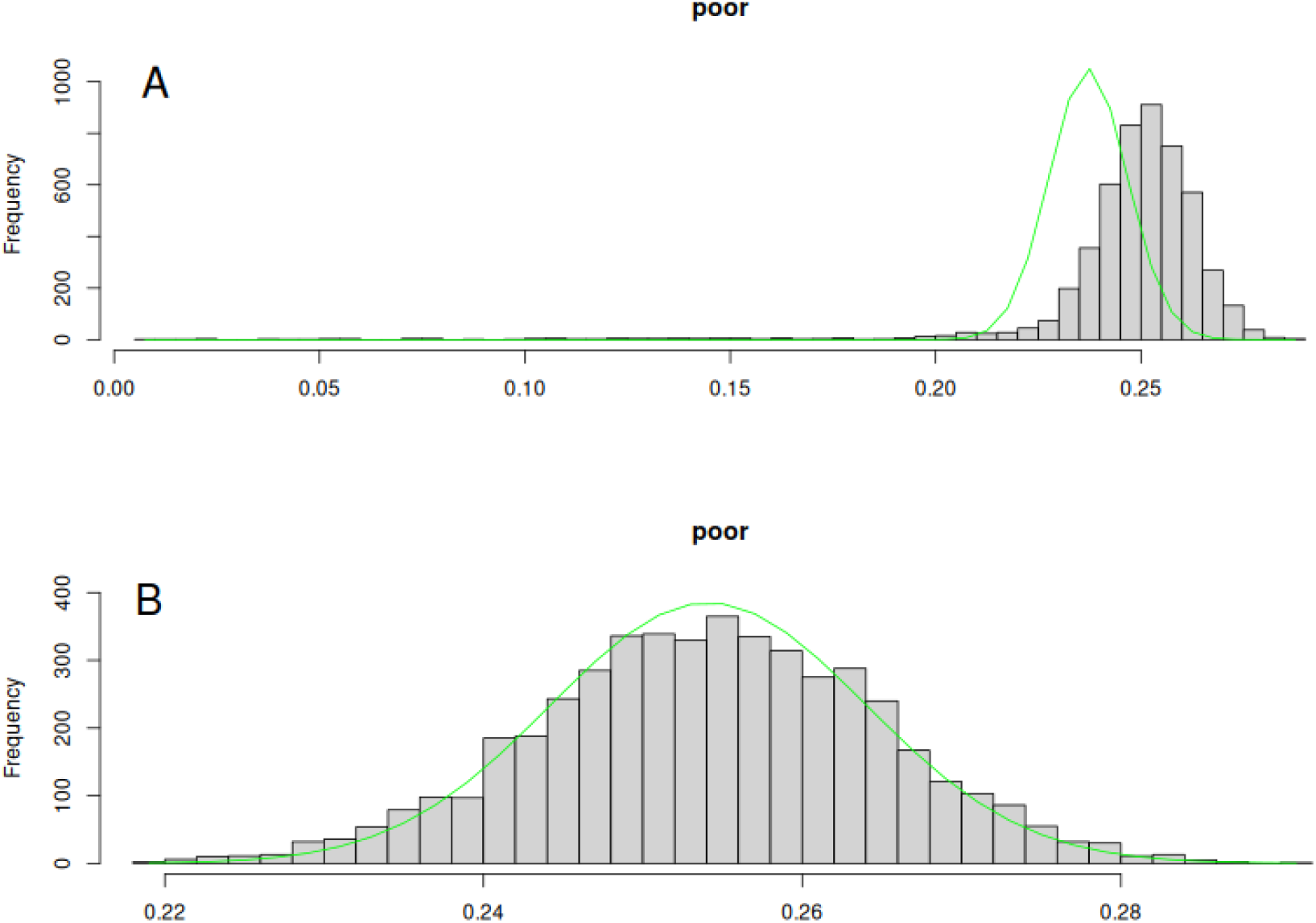
Checking for contaminated or (as in this case) degraded samples via het-erozygosity (`hetzninoo_fancy`). Example for school shark, as per Thomson et al. (2025). A: Preliminary, showing a long left-hand tail of degraded samples, and poor overall fit. B: A good fit after iteratively filtering outlier samples, filtering dubious loci based on `check6and4`, and re-estimating allele frequencies. (Note cleaning steps have been re-ordered from Thomson et al. to more clearly demonstrate `hetzninoo_fancy`.)

### 5.5. Consistency of kin-pairs

When aiming for HSPs, it is also worth checking internal consistency of the definite or likely HSPs, in terms of implied pedigree. For example, if sample A is allegedly a half-sibling to samples B, C, and D, then at least two of B, C, and D must be half- or full-siblings to each other^5^. However, we have encountered one situation where a handful of samples appeared to be half-siblings to large numbers of other samples that were clearly *not* siblings to each other. The cause turned out to be non-robustness of the PLOD statistic to rare null alleles at a few loci, whereby apparent but probably spurious double-nulls for the same locus in two animals led to a high likelihood of kinship. This motivated us to introduce the “3-way genotyping” option for loci with rare nulls (as in section 3.1), which prevents extreme values for single-locus LODs.

To help detect such situations, the function `chain_pairvise` shows inter-related clusters on a set of alleged HSPs— say, from `find_HSPs` with a threshold set deliberately lower than usual, to reduce the number missed. Note that `chain_pairvise` only makes sense when the potential 2KPs are very likely HSPs, as opposed to GGPs or FTPs, since the latter lead to quite different structures of inter-related groups. The kin-finding process is often made simpler by restricting 2KP comparisons to subsets where GGPs and FTPs are demographically unlikely.

Although these post hoc pedigree checks are useful for CKMR QC, we do not generally recommend using pedigrees to help find CKMR kin-pairs in the first place, because of the distorting effect on false-positive and false-negative rates (section 6.1). However, there are sometimes situations in CKMR where inter-related groups of HSPs and FSPs are inevitably frequent, for example when sampling larval fish. Kinship consistency then becomes particularly important for extensions of pairwise CKMR, requiring something more sophisticated than `chain_pairvise`. This is outside the scope of `kinference` per se, but the `sibgrouper` package (https://closekin.r-universe.dev/sibgrouper) provides tools to help.

## 6. Discussion

There are several ways to analyse kinships from genotypes, implemented in numerous programs that are subject to various assumptions and limitations, and are aimed at different objectives. Compared to other programs, `kinference` gives rather limited outputs and imposes rather restrictive assumptions, because of its specific context: data preparation for Close-Kin Mark-Recapture. This discussion provides our rationale, compares `kinference` to other software in a CKMR context, and explores future directions.

### 6.1. Likelihood, assignment, and pedigree

Unlike many other kin-finding programs, `kinference` deliberately does not make probabilistic assignments of pairs to kinship categories: there are no statements of the form “samples X and Y are most likely an FSP, with probability 99.7%”. Nor does it make claims of the form “samples X and Y are significantly different from Unrelatedness at *p* = 0.0001”, or attempt to link pairwise kinships to form pedigrees. Instead, it just reports pairwise pseudo-log-likelihood ratios from which the user can decide how to classify each pair, based largely on graphical outputs and perhaps taking into account other factors such as sample covariates and qualitative demographic knowledge. `kinference` also estimates the corresponding false-negative rate for classification of HSPs. As explained in section 2.2, these outputs from `kinference` fit naturally into a CKMR model, but it is worth examining why the above alternatives would be less suitable. The philosophical distinction between the different flavours of statements above are usually irrelevant since the kinship of most pairs is obvious, but there are major practical implications when it comes to borderline 2KPs.

A helpful description of different styles of “kinship assignment” is given in Weir et al. (2006). The most straightforward way to examine the kinship of a specific pair is by means of a likelihood ratio statistic: take two potential kinships (say, 2KP and UP), calculate the probability of the pairs’ observed genotypes under each, and take the ratio of probabilities of the two “hypotheses”. This can be very informative. Nevertheless, the likelihood ratio does not in itself allow any formal probability statement; for that, it would be necessary to introduce some kind of prior distribution on kinship, and to apply Bayes’ theorem, as Weir et al. (2006) describe. The context for prior-selection that they suggest, of identifying family remains after an airplane disaster, is about as far removed from CKMR as it is possible to imagine, and is a reminder of how diverse the applications and requirements of kin-finding can be.

Wang and Santure (2009) also emphasize the role of sibship and parentage priors in their section “The likelihood function” (although the “likelihood function” in their equation 1 does not conform to main-stream statistical usage). They suggest default choices in terms of parameters like “proportion of candidate parents sampled”. However, as noted in section 2.2, the whole point of CKMR is to *estimate* the very demographic parameters which would determine the priors on kinship, if used. Hence, it would be an obvious circular error to introduce such priors into kin-finding, which is merely the data-preparation phase for CKMR itself. `kinference` just reports and plots pairwise statistics from which the user can make final decisions. Since the scope of kin-finding is comparatively unambitious (POPs, FSPs, 2KPs), the appropriate decisions are usually pretty obvious given adequate data. The exception is when choosing a cutoff for retaining definite 2KPs, where `kinference` can propose a suitable cutoff based on a user-specified tolerance for false-positives from 3KPs and/or 4KPs; the corresponding false-negative rate is also estimated. This is “bounded-error classification”, rather than probabilistic assignment; no priors are involved.

The likelihood or likelihood-adjacent statistics calculated by `kinference` might momentarily suggest that formal significance tests or *p*-values are useful for CKMR kin-finding. They are not. The main problem is that a “null hypothesis” of unrelatedness in a kinship likelihood-ratio test is *a priori* far more probable than the alternative hypothesis of one specific kinship. With SBT, there are ∼ 10^8^ comparisons but only ∼ 10 POPs and HSPs, so that the appropriate *p*-value for any test would be far out into the tail, and hard to choose reliably. For other null and alternate hypotheses not involving UPs, the prior asymmetry may not be as strong but it would certainly influence Type-I and Type-II error rates. Of course, we do not want to get into the details of specifying demographic priors, but equally we cannot avoid the fact that they exist and are relevant to “significance”. The other problem is that the null hypothesis of unrelatedness is highly composite; there is really no such thing as an “unrelated” pair, just varying degress of kinship which are weaker than the alternative target kinship, and which have substantially different distributions for the test statistic. Further, unless the null hypothesis is UP or POP, linkage makes it impossible to fully predict the null/reference distribution (section 4.2.5). While *p*-values can be quite useful in CKMR QC (at least in a relative sense, for ranking), and near-tail probabilities are essential for estimating false-negative rates, in our experience formal significance tests and *p*-values at the level of individual pairwise comparisons have not been useful for CKMR kinship.

Another way to find close-kin is by constructing pedigrees (family links between several kin), as opposed to just pairs of kin. As noted by e.g. Jones and Wang (2010), pedigrees are certainly helpful when trying to find as many kin-pairs as possible (especially when pairwise statistics are ambiguous), and maximizing the information about the exact kinship of close relatives (for example, distinguishing different 2KPs). For CKMR, though, there are two problems. The first is practical: in large populations with sampling at the level required to find enough kin-pairs for reasonable precision, there will be few if any triads and beyond of close-kin. For example, with an adult population size in the millions, and a sample size such that 50-100 close-kin pairs are truly present, then only about 1% of the samples might be involved in any close-kin pairs, so quite probably there will be no triads at all amongst the pairs; looking for pedigrees would be pointless.

The second pedigree problem is more pernicious, and applies to smaller populations where kin-triads can be numerous. If pedigrees are taken into account, then HSPs in large families will be found more often, because a HSP whose kinship statistics happen to fall outside the target range can often be “rescued” based on other HSPs, FSPs, and POPs involving the pair-members. Such rescues cannot happen for HSPs which lack other sampled siblings. This would distort the false-negative probability and make it dependent on sampled family size, violating the assumption of constant false-negative rate that underlies equation 2.3.

Although we consider that pedigrees should be avoided when finding kin-pairs for CKMR, it can nevertheless be useful as a post hoc QC check amongst kin-pairs that have already been provisionally identified (Section 5.5).

### 6.2. kinference and alternatives

`kinference` provides a complete toolkit for going from genotypes to CKMR-ready pairwise kinships. Compared to many other relatedness programs, though, `kinference` imposes fairly strict assumptions (e.g. panmictic population) and data requirements (e.g. no missing values); see the Introduction. It produces comparatively unambitious output (e.g. pairwise kinship statistics, without pedigrees), and requires extensive user involvement rather than automating its decisions (e.g. which subsets to test for which kinships, in which sequence; acceptable tolerances; no defaults for parameters). All those decisions are motivated by top-down consideration of CKMR requirements.

Because of its limited scope and tight requirements, `kinference` offers the following key features:

1. Purely pairwise likelihood-based kinship statistics, with expected values for diagnostic purposes, for thousands of SNPs on tens-of-thousands of samples (section 4.2).
2. For HSPs, empirical allowance for linkage (no genome needed), with false-negative rate estimated as a consequence of user-specified tolerance for false-positive weaker kin (sections 2.2 and 4.2.5).
3. QC statistics for samples and loci, with accurate reference/null distributions for comparison (section 5).
4. Allowance for null alleles (section 3.1).

#1 and #2 are heavily entwined. For CKMR, it is much better to be “wrong” (i.e. to discard true HSPs) more often, but in a consistent and statistically predictable way, than it is to be be as “right” as possible for each potential kin-pair, at the cost of having variable and unestimatable error rates that may be linked to demographic structure (e.g. by constructing pedigrees). We regard #1, #2, and #3 as always essential for CKMR (unless the model will only use POPs, in which case #2 and probably #1 are irrelevant); and #4 is essential in many, though not all, CKMR projects.

Among many widely-cited programs that deal with relatedness, at least the following are all capable of identifying POPs, FSPs, and 2KPs: `HL-RELATE` (Kalinowski et al., 2006), `C0L0NY` (Jones and Wang, 2010), `KING` (Manichaikul et al., 2010), `sequoia` (Huisman, 2017), and `CKHRsin` (Anderson, 2024). Most can tackle some problems that are beyond the scope of `kinference`, such as population substructure, inbreeding, high genotyping error rates, and different types of genetic marker. However, none seem to offer all the CKMR-oriented features listed above. In particular, none (except `CKHRsin`, to some extent) directly address @2, and only `HL-RELATE` seems able to cope with null alleles. `COLONY` and `sequoia` are heavily pedigree-oriented, and `sequoia` especially uses demographic priors which, for CKMR, should be avoided. In the interests of being robust to population structure, `KING` sacrifices statistical efficiency by using non-likelihood-based statistics (rather than a PLOD) to find 2KPs; while this might be fine in the context of the GWAS-scale data (e.g. *>* 10^5^ SNPs) that `KING` expects, it is undesirable for the limited number of SNPs usually affordable in CKMR studies. `HL-RELATE` and `CKHRsin` compute basically similar PLODs to `kinference`. When it comes to converting PLODs into kinships, `HL-RELATE` simply assigns to each pair the kinship which maximizes the joint likelihood of the genotypes (actually a pseudo-likelihood, since linkage is ignored), which is not good for CKMR since it will lead to many false-positives; there is no option for user control of false-positive/negative issues.

Apart from `kinference`, the best option for CKMR is likely to be `CKHRsin`— unsurprisingly, given its name— which is the only other program to estimate false-positive/negative rates (albeit under an assumption of equal numbers of HSPs and HTPs, which is usually not the case), and which also acknowledges the existence and impact of linkage. However, `CKHRsin` is geared towards simulation and exploration, rather than direct data analysis. Its default estimates of false/positive rates assume no linkage, which would give misleading results for 2KPs (the main case where they matter). In order to adjust those false-positive/negative rates, the user has to specify linkage details in advance, something which would not have been possible in the CKMR applications we have seen.

Direct comparisons between these programs are difficult, because they are “tuned” for false-positive/ negatives in very different ways that can be opaque. Tsukahara et al. (2025) used real and simulated data to compare HSP-finding from `C0L0NY`, `CKHRsin`, and their own approach `fraRF`. They found substantial differences in the numbers of HSPs returned. This is not surprising in terms of the different tunings (e.g. default no-linkage assumption for `CKHRsin`; no options to control or to estimate false negative/positive rates for `C0L0NY`). They noted that, in their simulated data, `COLONY` and `fraRF` tended to produce roughly equal numbers of false-positives and false-negatives, so that the estimated number of HSPs was about right overall. However, this is not actually a good thing for CKMR, as 2.2 explains: the false-positive rate from HTPs to HSPs will be substantially higher for some covariate-strata than others, leading to bias in parameter estimates. From a modelling perspective, it is much better to keep the overall false-positive rate low, and to estimate the (much higher) false-negative rate reliably, and then allow for the latter within the CKMR model. The Tsukahara et al. paper illustrates well both the importance of doing so, and the difficulty of doing so with other software.

### 6.3. Future directions

`kinference` has been working well for CKMR projects since its first application to SBT around 2017. It has kept pace with modest improvements in genotyping, both in sequencing and in other lower-cost targeted assays. However, genetics has moved rapidly, and there are several op-tions for extending `kinference` to take advantage of affordable improvements available now in 2026, without even considering newer approaches such as long-read sequencing.

#### Computing speed

As genotyping costs drop and sampling logistics improve, CKMR projects are growing in size. At present, a routine like `find_HSPs` takes just a minute or so for around 15,000 samples (∼ 10^8^ pairs) with a few thousand loci. However, planned projects with ∼ 10^5^ samples might entail ∼ 10 pairwise comparisons, so speed might become an issue. The PLOD-calculating routines such as `find_HSPs` are already reasonably efficient simply because they are written in C++. However, unlike KING for example, `kinference` does not take advantage of modern features like multicore parallelization, SIMD, or GPUs, whereby substantial speed gains could be made.

#### Microhaplotypes

A properly-genotyped microhaplotype (for example, several SNPs within the ∼ 100bp sequence-length of one locus, or a triallelic single SNP) undoubtedly has more kin-finding power than the same locus “reduced” to biallelism by aggregating alleles into just two categories. Thus, microhaplotypes have been suggested as a way to increase kin-finding power without costing more (Baetscher et al., 2018). In our experience, they have been quite common in tuna, but much less so in other species. In principle, the `kinference` calculations could be extended fairly easily to microhaplotypes, but there are practical issues with the code; and our early experiences with trying to call microhaplotypes from sequence counts were not encouraging. Although microhaplotypes could be useful in some applications, we are skeptical that the work required to extend `kinference` would be worthwhile in general, especially compared to the next option.

#### Genome-assisted kin-finding

Historically, high-quality de novo genome assemblies were unavailable or unaffordable for most species, but that has changed since the early 2020s. As of 2026, a chromosome-level genome assembly for a new species is often on the same order of cost as the pilot RAD-based sequencing needed to find kinship loci, and is therefore a small fraction of the total budget of a big CKMR project. By knowing the location of each kinship locus within each chromosome, the statistical power of kin-finding can be greatly increased. For example, Ramstetter et al. (2017) report useful accuracy down to at least 4KPs, admittedly using essentially perfect data for humans; the extent of benefit for other species will vary, depending on chromosome counts, crossover rates, and number of markers. For CKMR, genome assemblies could be “retro-fitted” to existing studies without needing extra loci or changes to genotyping per se. This should reduce, and for some species perhaps even eliminate, the false-negative/positive annoyance with 2KPs. However, the algorithms needed for genome-assisted kin-finding are very different to the comparatively simple PLOD calculations in `kinference` and similar software. They have generally been developed for hyper-dense and well-understood marker sets on humans (`KING` oflers one option); redesign would be required for wild-animal CKMR settings with far fewer SNPs. Even when high-quality genome assemblies are available, though, we expect that the existing routines in `kinference` will continue to be important for QC, POPs and FSPs, and initial screening of 2KPs to remove billions of clearly-uninteresting UPs.

## Acknowledgements

CKMR would not be where it is without Diversity Arrays Technology (DArT), who produced all the high-quality SNP data used for kin-finding at CSIRO. David Lawrence Miller wrote the first version of the C++ code. Luke Lloyd-Jones gave a very helpful review of the manuscript. The development of `kinference`, and the generation of data used with permission in this manuscript, was supported by CSIRO alongside various organizations in Australia and overseas: FRDC, AFMA, CCSBT, and the Department of the Environment.

## Appendix A. Saddlepoint Approximations to null/reference distributions

Saddlepoint approximations (SPAs) are widely used in statistics to approximate probability densities and cumulative probabilities of some statistic *X* (usually one-dimensional). In general, they are highly accurate even in the tails of the distribution, and they do not require simulation (Barndorff-Nielsen and Cox, 1989, section 4.3; Butler, 2007). To implement a SPA, the moment generating function (MGF) 𝔼 [exp (*tX*)] must be computable. This is conveniently true for several statistics in `kinference` that comprise sums across independent loci, so that the overall MGF is the product of per-locus MGFs, and the per-locus expectation is merely a weighted sum over the set of possible genotypes at that locus. Here we explain a few computational aspects common to all SPAs in `kinference`; the actual MGF for each statistic is given in the appropriate section.

The basis for a SPA is not the MGF per se but its logarithm— the cumulant generating function *J* (*t*)— and the first two derivatives thereof. These automatically satisfy *J* (0) = 0, *J*′ (0) = 𝔼 [*X*], *J*″ (0) = 𝕍 [*X*]. The probability density^6^ *f* (*x*) ≜ |*d* ℙ [*X < ξ*] */dξ*|_*ξ*=*x*_ by has a first-order SPA 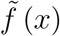 defined

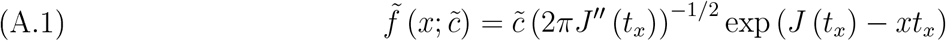

where *t*_*x*_ satisfies *J*′ (*t*_*x*_) = *x*, and 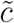 is a normalizing factor discussed shortly, taken as 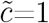 if omitted. If the SPA is used to compute 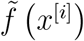 for specific values *x*^[*i*]^, then each *t*_*x*[*i*]_ requires numerical solution. But this somewhat slow and tedious step can be avoided when we want to approximate the entire density *f* (); instead, we can take a grid over a range of *t*^[*i*]^ (wide enough to cover the tails) and then, for each one, directly calculate the corresponding *x*, i.e. *x*^[*i*]^ = *J*′ (*t*^[*i*]^). These can be plugged into eqn (A.1) and used to fit an interpolating spline through the pairs 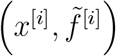.

This approach also helps with the normalizing factor. While 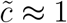 is a reasonable approximation in its own right, better SPA performance is usually obtained by ensuring that the SPA integrates to 1 across the range of *X*, i.e. that 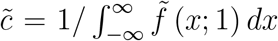. Making the change-of-variables *x* ⟼ *t*_*x*_ and writing *t* instead of *t*_*x*_ for short, we have

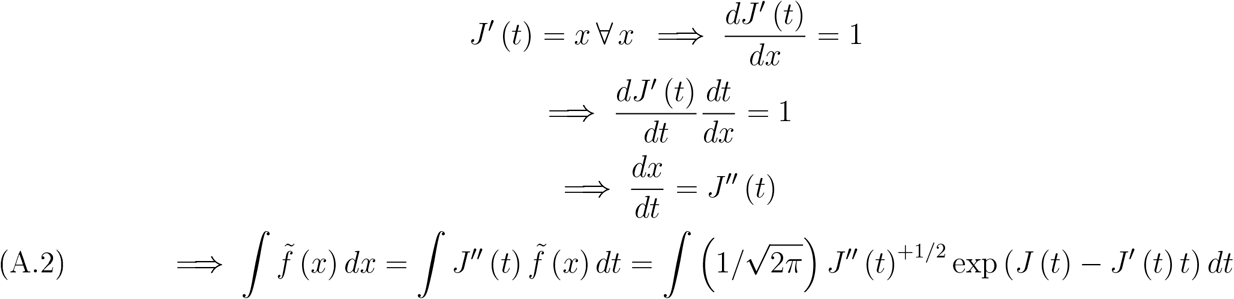

The final term in eqn (A.2) can be approximated by summing across the grid *t*^[*i*]^, so we can obtain an improved normalizing factor 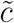 with minimal work.

The remaining issue is what range of *t* to cover with the grid, say between ±*t*^∗^. This also defines the effective *X*-range, since *x*^*[i]*^ = *J′*(*t*^[*i*]^. We start with the approximation *J*′ (*t*) ≈ *J*″ (0) + *tJ*″ (0) = 𝔼 [*X*] + *t*𝕍 [*X*]. Pretending for the moment that *X* is Gaussian, if we require *t*^∗^ to cover (say) a range of *X* out to 6 standard deviations from its mean, then we would need 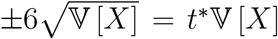 so that 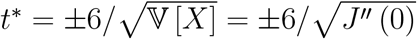. Of course, *X* is not actually Gaussian (otherwise we would not need a SPA), but 6-10 standard deviations should be ample to cover the relevant tails regardless, and numerical difficulties can result if the range of *t* is made too large.

The above calculations feature in several functions of `kinference` and the supporting package `gbasics`, including `prepare_PL0D_SPA` which can be inspected for further details.

Finally, we note that there are alternative forms of the SPA for direct calculation of tail probabilities (e.g. the “Lugananni-Rice approximation”; Barndorfl-Nielsen and Cox, 1989, equation 4.71). However, in our experience, simpler and more numerically reliable code comes from using the above density approximation, and integrating where necessary to obtain interval or cumulative probabilities.

## Appendix B. Accelerated EM algorithm for allele frequencies

Given *n* samples, if the 6-way counts (i.e., of AA, AO, AB, BB, BO, and OO genotypes) were known accurately, then the MLE of *π*_B_ would be (2*n*_BB_ + *n*_BO_ + *n*_AB_) */*2*n* and similarly for *π*_*O*_. This is the M-step of the EM algorithm. The unobserved sufficient statistics for *π*^*^ are therefore *n*_BO_ and *n*_AO_, given that *n*_BBO_ = *n*_BO_ + *n*_BB_ etc., and that the 4-way genotyping totals are known. In the E-step, the conditional expectations of the unobserved sufficient statistics, given working estimates *π*^∗^, are computed from

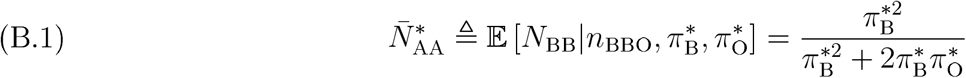

and similarly for *N*_AA_, *mutatis mutandum*. The conditional expectations are then used in place of the unobserved true values in the next M-step, and so on; this is the EM algorithm described in Dempster et al. (1977). Starting from any approximate estimate, such as 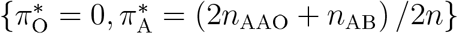, it will ultimately converge to the MLE; the convergence rate per iteration is much slower than direct MLE, though each iteration of EM may be much faster, as in this case. We speed up convergence by applying an Aitken acceleration (e.g. Press et al., 1992, section 5.1) after every few steps of EM. Specifically, we take the recent sequences of 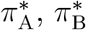, and 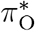, extrapolate each one to where it might converge, then rescale the three extrapolands to sum to unity. In most cases, convergence to machine precision is achieved within 10 Aitken steps.

## Appendix C. Optimal weights for pseudo-exclusion

For any candidate kin-pair, our pseudo-exclusion statistic in section 4.4 is the weighted sum across loci of the number of pseudo-exclusions, i.e.

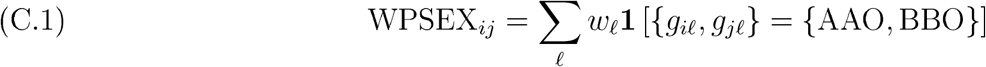

The optimal choice of weights will give the best separation between the two distributions of WPSEX for true UPs and for true POPs. Specifically, we seek the weight vector *w*^∗^ which maximizes the number of standard deviations between the expected values for POPs and UPs, or

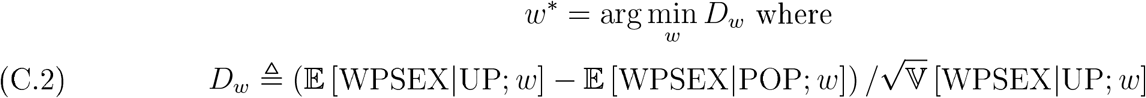

Loci with higher null frequency will receive lower weights, because they are more likely to give a pseudo-exclusion due to coinheritance of a null. Since any locus in a UP can generate a pseudo-exclusion simply by chance, the weights are not heavily skewed towards particular loci.

To derive the optimal *w*^∗^, let *p*_0*ℓ*_ and *p*_1*ℓ*_ be the pseudo-exclusion probabilities at a locus for UPs and POPs respectively, i.e.

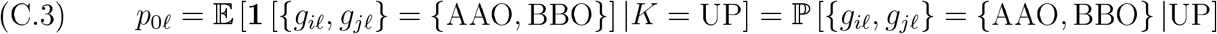

Omitting *ℓ* for brevity, at any single locus with allele frequencies (*π*_A_, *π*_B_, *π*_O_), elementary calculations give

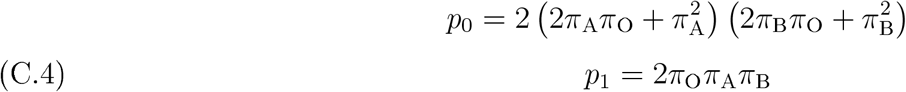

We first prove the intuitively obvious result that pseudo-exclusions are always more likely for UPs than for POPs, i.e. that *p*_0_ *> p*_1_. The ratio is given by

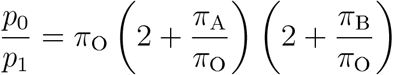

For any given *π*_O_, the sum *π*_A_ + *π*_B_ ≡ 1 − *π*_O_ is fixed, so the product of the parenthesized terms is minimized when either *π*_A_ = 0 or *π*_B_ = 0. Since that is a minimum, we have

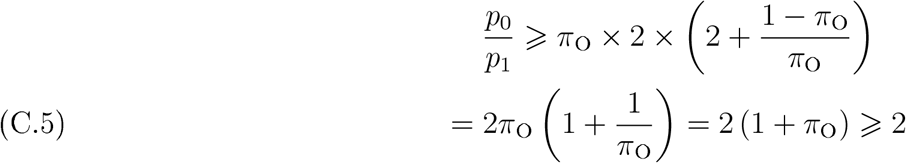

so that exclusions are always at least twice as likely for a UP than for a POP. It follows that *D*_*w*_ *>* 0.

Then, since loci are independent for both UPs and POPs, we can define the following:

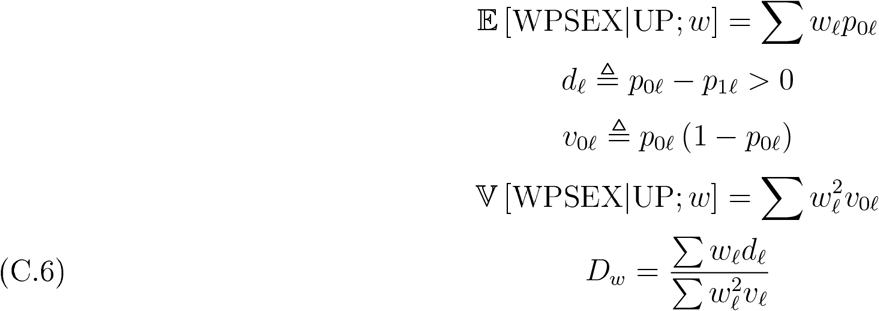

To find the optimum *w*^∗^, note first that *D*_*w*_ is unchanged when the whole vector *w* is multiplied by any positive constant. We can therefore solve for *w*^∗^ subject to any convenient scaling constraint, and then rescale *w*^∗^ in whatever way we choose. It is quite tedious to work directly with the standard constraint of Σ *w*_*ℓ*_ = 1, but the solution is readily found if we work instead with the constraint 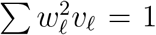, so that *D*_*w*_ = *w*_*ℓ*_*d*_*ℓ*_ when the constraint is satisfied. Using a Lagrange multiplier *λ*, we therefore need to solve

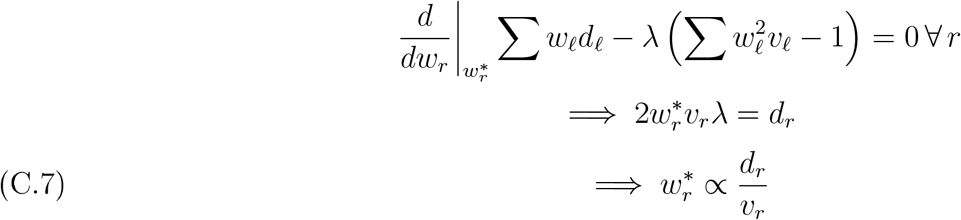

Since *d*_*r*_ ⩾ 0 and *v*_*r*_ ⩾ 0, these optimal weights can all be chosen positive.

## Appendix D. Optimal weights for qc checks

The per-sample “hetzminoo” QC statistic *H*(*w*) = Σ_*ℓ*_ *w*_*ℓ*_*S*_*ℓ*_ of section 5.4 can be tuned by picking the locus-specific weights *w* for high power, to detect either dropout (usually from degraded DNA) or contamination. We first consider the general problem of choosing weights to detect “badness”. Suppose that “badness” of sample *i* can be parametrized by some scalar *θ*_*i*_; the meaning of *θ* will differ between contamination or dropout, leading to different explicit formulae for *e*_*ℓ*_ and *v*_*ℓ*_ (the mean and variance of *S*_*ℓ*_) in terms of *θ*, as explained later. For high statistical power, we want the expected value of *H* (*w*) for a bad sample with *θ* = *θ*^∗^ ≠ 0, say, to be far out in the tail of the null distribution for good samples, i.e. many standard deviations away from the expected value when *θ* = 0. In mathematical terms, we want to choose *w* so that the criterion 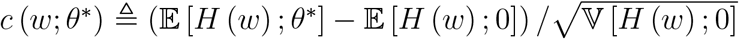 is as large in magnitude as possible^7^; the sign of “large” depends on the type of badness.

The loci are independent by assumption, so 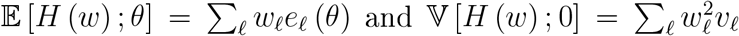 (with the convention that the argument to *e*_*ℓ*_ or *v*_*ℓ*_ is 0 if omitted). Now, *c* (*w*; *θ*^∗^) depends on *θ*^∗^ which is unknown, but our main interest is in small *θ*^∗^ since samples with large *θ*^∗^ should stand out clearly. As *θ*^∗^ → 0, clearly *c* (*w*; *θ*^∗^) → 0 also, but we can take a Taylor expansion:

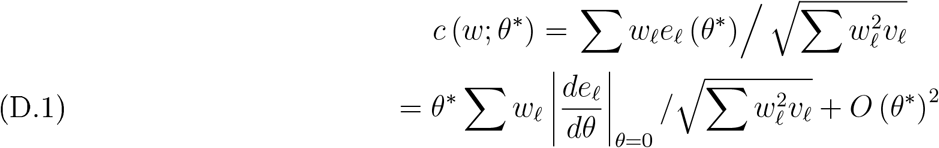

Because *θ*^∗^ appears only as an overall multiplier, the optimal weights do not depend on the extent of badness, provided that badness is small. Defining *δ*_*ℓ*_ ≜ |*de*_*ℓ*_*/dθ*|_*θ*=0_ (with explicit formulae given later), equation (D.1) then has the same form as equation (C.6) for the WPSEX *D* (*w*). The same Lagrange-muliplier approach leads to optimal weights 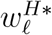 satisfying

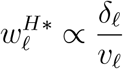

These can be multiplied by any constant to achieve whatever constraint is required, say 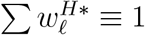.

To compute *δ*_*ℓ*_ and *v*_*ℓ*_, first consider a good sample without contamination or dropout, i.e. *θ* = 0. The scored genotype is the same as the true genotype, so that

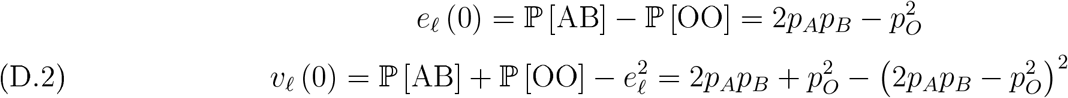

where genotypes without the qualifier “scored” refer to the true genotype of the target sample, and {*p*_*A*_, *p*_*B*_, *p*_*O*_} are again the major, minor, and null allele frequencies.

We now need to define suitable models and *θ*-measures for the two types of badness, leading to formulae for ℙ [AB scored] and ℙ [OO scored] in terms of *θ* and the minor- and null-allele frequencies. From those, we can calculate *e*_*ℓ*_ (*θ*; contam) and *δ*_*ℓ*_ (*θ*; contam), and the optimal weights 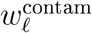 and 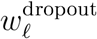. The models do not need to be perfect representations of badness; their role is to help choose good weights for detecting badness in individual samples. Although the two sets of weights can be reasonably different, in practice we have seen little difference between the variants, in that the same samples tend to show up as outliers regardless of variant. Nevertheless, on theoretical grounds we would recommend looking for high-*H* outliers with the contaminated version, and for low-*H* outliers with the dropout version.

### D.1. Contamination

For contamination, assume that a proportion *θ* of loci in the target sample have their genotypes perturbed by the addition of a genotype from a contaminator, to the extent that any allele present in either sample will be scored. If the target sample is truly a heterozygote, then the scored genotype will also be a heterozygote regardless of any contamination. If not, then a heterozygote can still be scored if the combined target and contaminant contain both alleles:

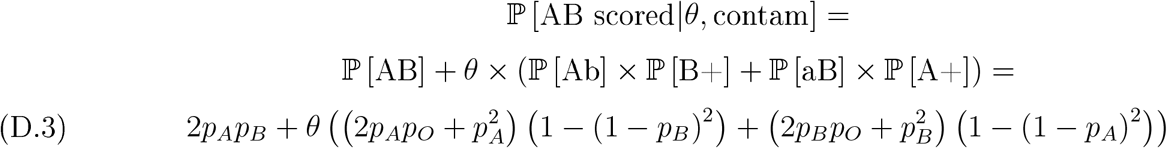

where the notation ℙ [Ab] means “at least one A and no B” (note the lower-case), and ℙ [B+] means for “at least one B”, etc.; those can all be expressed in terms of underlying allele frequencies. To score a double-null, the target locus must be truly double-null, and the contaminant (if any) must be as well. Thus:

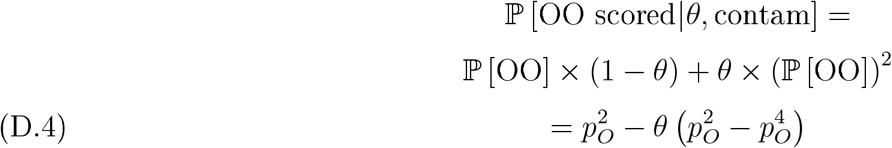

Thus

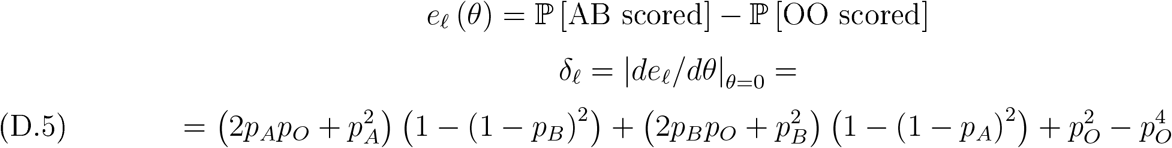

### D.2. Dropout

We assume that dropout can cause one copy of an allele to be missed with probability *θ*, but that it is unlikely to miss two copies. (If dropout is severe, it will aflect many loci, and be clearly detectable; we are focusing on low levels of dropout.) Thus:

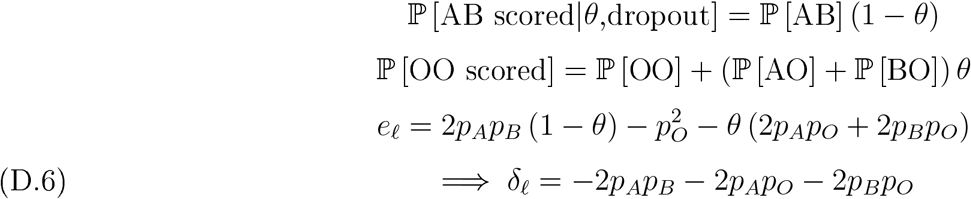

### D.3. Null/reference distribution

It is useful to know what the null/reference distribution of *H* should look like for “good” samples, i.e. *H* (*w*^contam^; *θ* = 0) and *H w*^dropout^; *θ* = 0 . The distributions turn out to be appreciably non-Normal, but since the loci are independent we can readily derive a SPA, which are known to give accurate tail probabilities (e.g. Butler, 2007). The approach is the same for the contamination and dropout versions (although the values will difler), because we are only interested in distributions when *θ* = 0 and the genotypes are in HWE.

The basis for the SPA is the moment generating function for a single locus *ℓ* (mostly omitted for brevity), i.e. 𝔼 [exp (*twS*)] for any value of *t*. Since *S* can only take the values {−1, 0, +1}, this is straightforward:

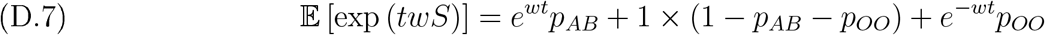

where for brevity we use e.g. *p*_*AB*_ instead of ℙ [AB], etc. Now defining *q* ≜ 1 − *p*_*AB*_ − *p*_*OO*_, we can derive the cumulant generating function *J* (*t*) and its first two derivatives:

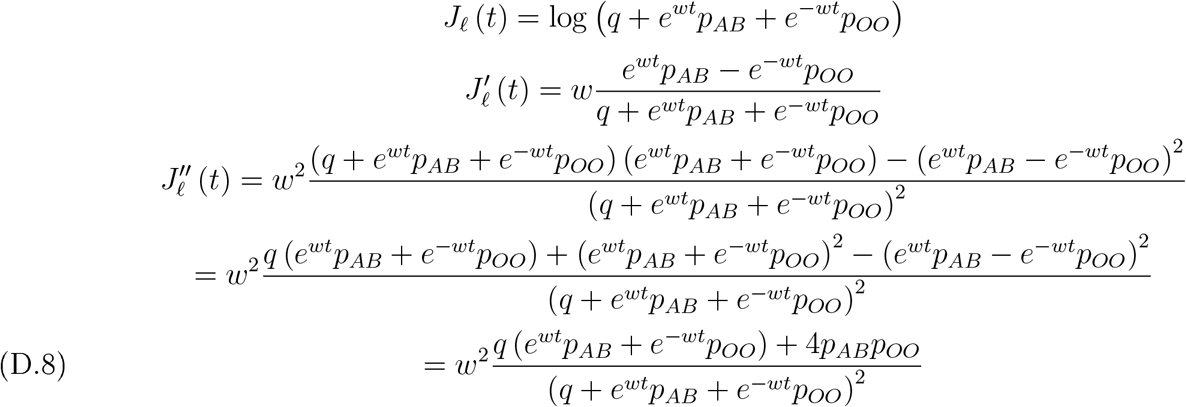

By independence of loci, the cumulant generating function of *H* is *J* (*t*) = Σ_*ℓ*_ *J*_*ℓ*_ (*t*), and similarly for the derivatives. The null/reference distribution of “hetzminoo” can then be obtained by SPA (Appendix A).

## Appendix E. Variance bounds for weaker kin

To set bounds on the variance of PLOD_HSP,UP_ (or just “PLOD” for short) for HTPs and HCPs, we consider the two most extreme ways that linkage might work:

1. there are *C* similar chromosomes, each containing a proportionate mix of all kinship loci, with no recombination;
2. there is just one chromosome, with kinship loci equally spaced and in random order, and with a constant and independent recombination rate *ρ* between loci.

For each case, we can predict the PLOD variance for any single-lineage kinship parametrized by *κ*_1_, as a function of *C* or *ρ*. Thus, we first estimate *Ĉ* or 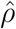 by matching the empirical variance of the right-hand size of the HSP bump where 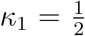, and then plug that estimate into the same formula when 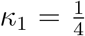 and 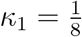 to predict the HTP and HCP variances. (For case 2, though not case 1, the specific type of kinship matters, as well as the overall order, so in this section we refer specifically to HTPs rather than 3KPs, and so on.)

The key formula is the Law Of Total Variance:

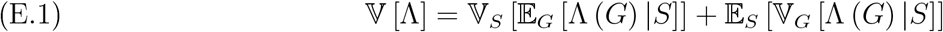

where *G* is the multilocus genotype, Λ (*G*) is the PLOD constructed as a sum of per-locus LODs *Σ λ*_*ℓ*_ (*G*_*ℓ*_), and *S* is the vector of coinheritance status across all *L* loci, with *S*_*ℓ*_ = **1** [coinheritance at locus *ℓ*]. For brevity, we mostly omit the argument *G*, and note that the distribution of *S* is implicitly conditional on kinship. Some further notation will be needed:

- [.|1] and [.|0] means “event Dot at a locus, given that the locus (1) is coinherited, or (0) is not coinherited”.
- The *aierape* per-locus means, moments, and variances of the LODs are 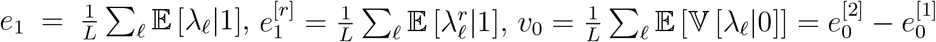, etc.
- *p*_1_ is the marginal probability of coinheritance at any locus. For single-lineage kin (i.e. not FSPs or descendents), *p*_1_ = *κ*_1_ = 2^−*m*^ where *m* is the number of meioses separating the two kin, so 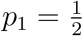 for HSPs (and GGPs), 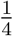 for HTPs, etc. Also let *p*_0_ = 1 − *p*_1_.

There is an implicit assumption below that the available loci are homogenously distributed across the many chromosomes or along the one chromosome. Since there are sure to be a few thousand loci, the Law Of Large Numbers provides some reassurance.

### E.1. Many chromosomes, no recombination

The genome is divided into *C* chromosomes each with *L/C* loci, and there is no crossover within a chromosome. Thus, for HSPs, the number *S* of chromosomes that an animal coinherits with its half-sib is Binomial (*C*, 1*/*2) and for a HTP it is Binomial (*C*, 1*/*4), etc. For

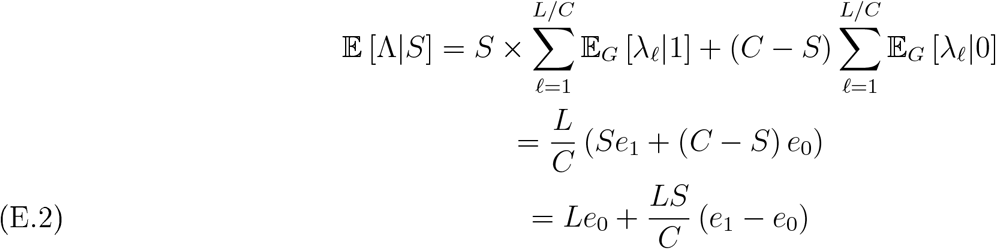

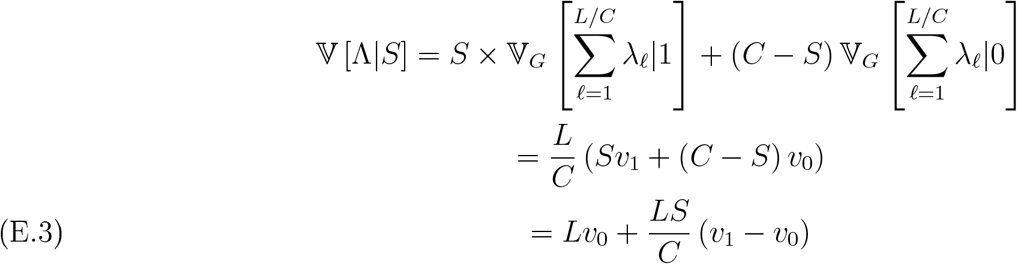

Plugging in the Binomial moments for *S*, for single-lineage kinship *κ*_1_ we get:

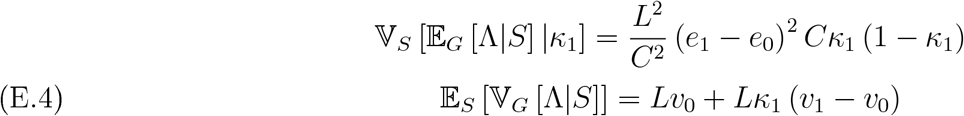

### E.2. One chromosome, frequent recombination

Unlike case A, it is now not just the kinship-order that matters, but also the type of kinship within order; for example, HSPs are different from GGPs. `kinference` only considers the commonest CKMR situation of “generalized half-cousins” HxCyP, which descend from a HSP with “x” additional meioses in one limb and “y” in the other, as in Figure E.1. Thus, H0C0P ≡ HSP, H1C0P ≡ HTP, and H1C1P ≡ HCP.

For HxCyPs, there are multiple coinheritance stages to consider: the initial half-sibling stage, plus the zero or more meioses within the left- and right-hand direct descent legs. There is no inbreeding, so an allele is coinherited in the green samples if-and-only-if it is coinherited at every stage.

**Figure E.1.**
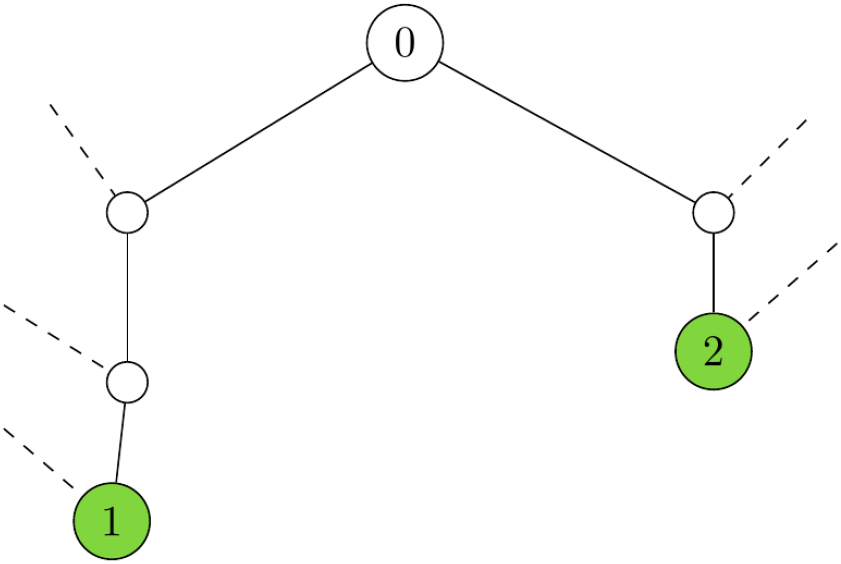
Coinheritance pattern for H2C0P (or H0C2P). Dashed lines indicate un-shared ancestors; unsampled ancestors are white circles; samples are green circles. Samples 1 and 2 share unsampled ancestor 0.

The mathematical approach here is similar to Hill and Weir (2011), who also address more complex situations. We assume that recombination at each meiosis follows a continuous Markov process with rate *ρ* per unit distance between loci (“Haldane’s mapping function”; Haldane, 1919). The probability that a recombination occurs between two loci a distance *d* apart is 1 − exp (−*ρd*), and the expected number of such recombinations is *ρd*. (The units of distance are irrelevant since we can scale *ρ* accordingly, so we may as well imagine ordering the loci along the genome/chromosome, and just using the index of each kinship locus from 1 to *L* to measure “location”.) First consider a single meiosis where 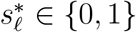 indicates inheritance of the “correct” copy at locus *ℓ*. If 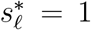, then the correct copy will also be inherited at locus *ℓ* + *d* (i.e. 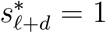) if-and-only-if the number of intervening recombinations is even, i.e. with probability 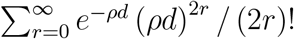. By comparison with the series expansions of cosh *x*, it can be shown that

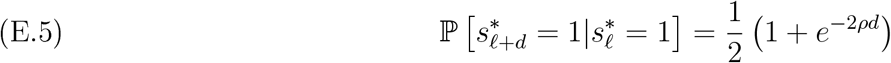

Note that the effective recombination rate in the first (HSP) stage is actually 2*ρ*, because crossover could happen in either of the two meioses. Because all stages have to coinherit, the overall conditional probability of coinheritance is

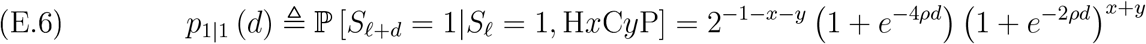

Since the marginal coinheritance probability is *p*_1_ for all loci, we can use the overall balance equation for a Markov chain to calculate *p*_1|0_(*d*) from the following (omitting the *d*-argument for brevity):

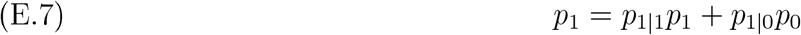

whence we can also calculate *p*_0|1_ = 1 − *p*_1|1_ and *p*_0|0_ = 1 − *p*_1|0_. Also, the detailed balance equation gives *p*_1|0_*p*_0_ = *p*_0|1_*p*_1_.

Writing *λ*_*ℓ*_ instead of *λ*_*ℓ*_ (*G*_*ℓ*_) for brevity, the PLOD Λ can conveniently be written as a function of coinheritance status *S* and its complement 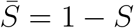 as:

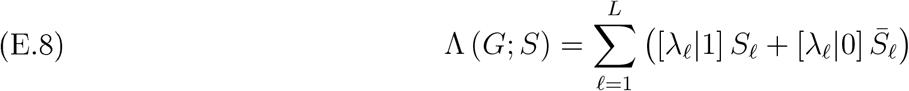

so that

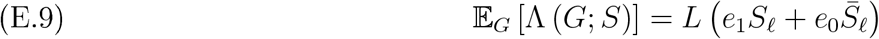

Since 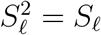 and 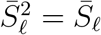, we also have

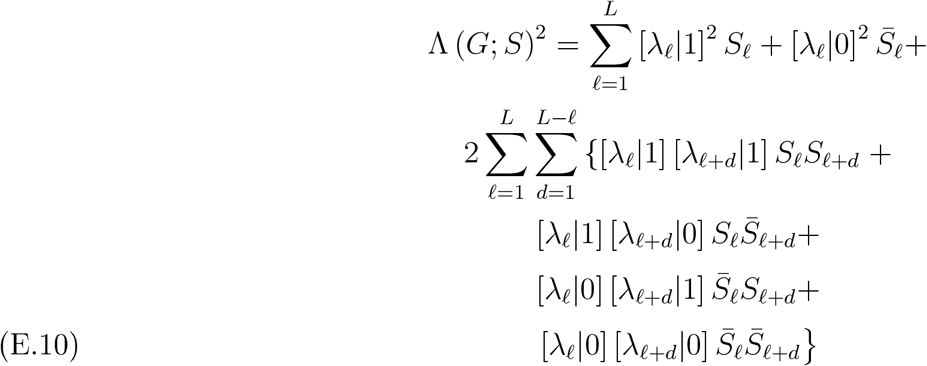

Making use of the detailed-balance and symmetry properties of the Markov process, this gives

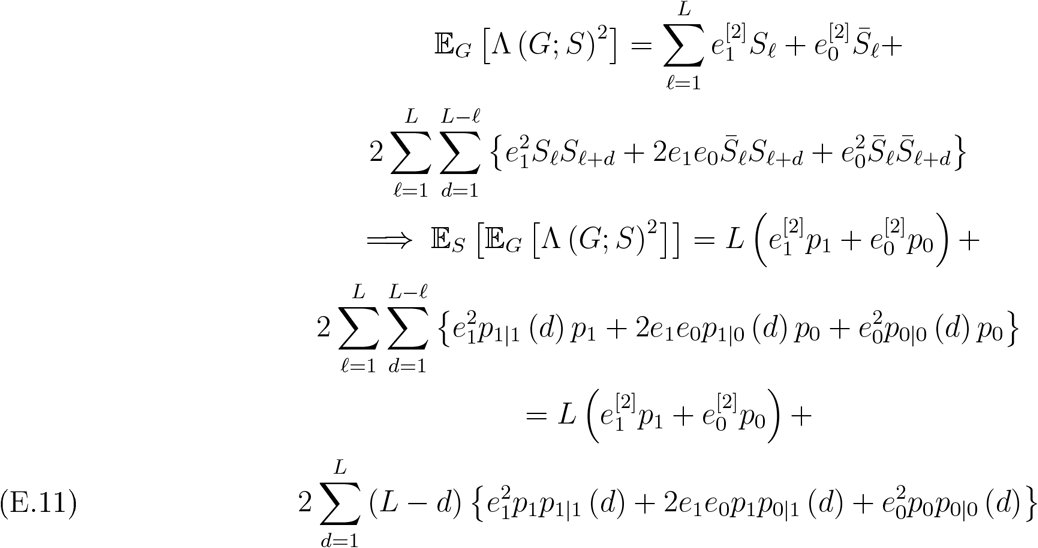

Finally, the left-hand term in (E.1) follows by substituting (E.11) and (E.9) into the identity 𝕍_*S*_ [𝔼_*G*_ [Λ]] ≡ 𝔼_*S*_ 𝔼_*G*_ [Λ] −(𝔼_*S*_ [𝔼_*G*_ [Λ]]) . The right-hand term in (E.1, 𝔼_*S*_ [𝕍_*G*_ [Λ]], is the same as in (E.4), since the marginal distribution of *S*_*ℓ*_ is the same under both linkage cases.

Although the sum in (E.11) can be re-written in terms of geometric sums and their derivatives, by substituting the values *p*_.|._ from (E.6), it is simpler and less error-prone to let the computer do the summation.

## Appendix F. Null alleles and Ö-way genotyping

In 2017 when the SNP version of SBT CKMR was being set up, it became apparent that null alleles were unavoidably common in SBT. Given the state of sequencing technology at the time, the most cost-effective approach was to keep the number of loci fairly low, but deliberately increase the read-depth so that single-nulls could be reasonably well distinguished from true homozygotes, where the read-depth of the non-null allele would be twice as high on average. For a locus where nulls are fairly common and the discrimination is reasonably good, this effectively adds a 3rd allele, substantially increasing the kin-finding power.

This paper does not generally deal with calling of genotypes, but 6-way genotypes require special treat-ment in `kinference` and it is worth explaining why. Figure F.1 shows the distribution of total read-depth for one well-behaved locus across several thousand SBT samples. The right-hand bump comprises het-erozygotes as well as true homozygotes; the central bump comprises single-nulls; and the spike on the left consists of double-nulls. The solid vertical lines show where the genotype-calling software (internal CSIRO code) makes its calls; if a sample is not a heterozygote, and not a double-null (i.e. to the right of the purple line), then it will be called as a single null if its read-depth is to the left of the turquoise line, or a homozygote if to the right. Although the separation between the central and right-hand bumps is fairly good, so that most single-nulls will be called correctly, there will be a significant proportion of errors between single-nulls and homozygotes. The genotype-calling software estimates that error-rate based on the fitted Normal curves, as shown in the figure; the turquoise line is placed so as to equalize the error rate in each direction. Within `kinference`, those error rates are used to turn the HWE-based probabilities of true genotypes into probabilities of *called* genotypes, conditional on the number of coinherited alleles. The code is fairly complicated.

**Figure F.1.**
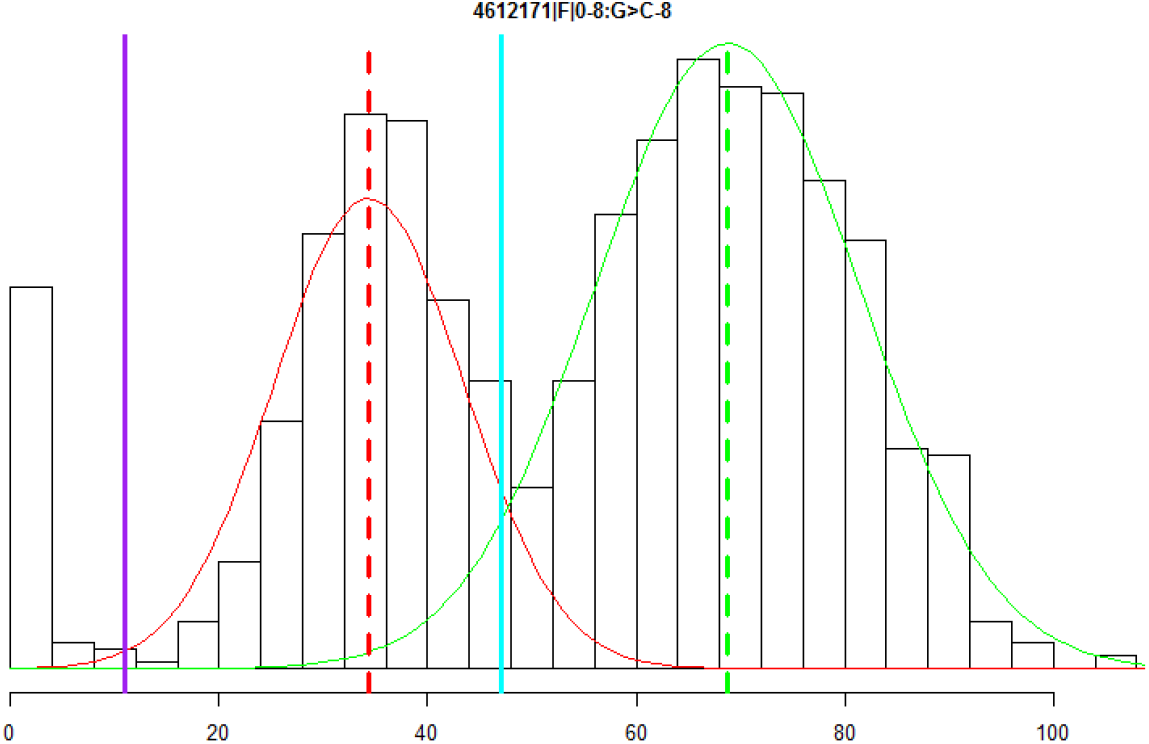
6-way genotyping for a well-behaved locus. The histogram shows the distribution across samples of normalized combined read depth for both non-null alleles. Solid vertical lines are cut-points for calling genotypes; dashed vertical lines are mean read-depths for heterozygotes and single-nulls; curves are fitted Normal distributions.

It is important to check whether 6-way genotypes at each locus are consistent with HWE, so the `check6and4` routine also plots observed and expected numbers for each 6-way genotype, allowing for the specific genotyping errors above. When the bump separation at a locus is not as good as in Figure F.1—for example, because average read-depths are lower at that locus— then it may fail HWE at the 6-way level, but still work perfectly well as a 4-way locus where AO and AA calls are aggregated into AAO. Overall, about 68% of the (roughly 1500 total) SBT loci are used at 6-way level.

This approach has worked reasonably well for SBT, in the sense that the results seem reliable and the total number of loci required for satisfactory identification of HSPs is considerably less than if only 4-way genotyping was used; back in 2017, that led to a significant cost saving, despite the higher read-depths required. On the other hand, 6-way genotyping is rather sensitive to small technical changes in the genotyping process; for example, if average read-depth changes, then so does the extent of overlap in Figure F.1. Considerable eflort has been needed to maintain consistency across nearly a decade of SNP genotyping. Nowadays it is more robust and efficient to simply use more loci in the first place, and to stay with 4-way (or 3-way) genotyping.

## Footnotes

1 Throughout, we use the term null/reference distribution in place of the more usual null distribution, to minimize confusion with null alleles.

2 This does not completely specify kinship, e.g. by not distinguishing GGPs and HSPs, but the latter is impossible using only the inputs to `kinference`.

3 To improve linguistic ow, we use “coinheritance” as a synonym for “being ibd”, and so on.

4 In general, likelihood ratios are optimal test statistics between two non-composite hypotheses; this is the Neymann-Pearson lemma.

5 B shares either mother or father with A. If C is not a half-sibling to B, then it shares either father or mother with A, respectively. If D is not a half-sibling to B, then it must share the same parent with A as C does.

6 Strictly, if *X* has finite support (which is true for all `kinference` uses), the probability density function is discontinuous, but the SPA typically still works well for approximating *integrals* of the density (i.e. interval probabilities) over small or large ranges of *X*; see Barndorff-Nielsen and Cox, 1989, chapter 4.

7 Actually it is debatable whether we should focus on the tail of the null distribution or on the tail of the alternative distribution when *θ* = *θ*^∗^, i.e. on whether the variance should be with respect to *θ* = 0 or *θ* = *θ*^∗^. However, the distinction vanishes anyway in the limit as *θ*^∗^ → 0.

## Notes

### Competing Interest Statement

The authors have declared no competing interest.

